# A unified framework reveals how evolutionary scale and abundance weighting shape phylogenetic community structure

**DOI:** 10.64898/2026.10.07.757245

**Authors:** Oskar Modin, Frank Persson

## Abstract

Metrics describing phylogenetic community structure are widely used to infer assembly processes in biological communities, but existing metrics capture only fixed phylogenetic scales and provide limited control over how rare and abundant taxa influence ecological patterns. Here we introduce *q*-weighted interpolated mean phylogenetic distance (^q^iMPD), a framework that continuously interpolates between broad evolutionary divergence and nearest-taxon distance while incorporating tunable abundance weighting. Applied to 3661 wastewater microbiomes, ^q^iMPD revealed that abundance weighting had little effect on broad-scale phylogenetic dispersion but strongly altered local phylogenetic structure. Null-model analyses showed that within-community signals of phylogenetic overdispersion and clustering, and between-community signals of phylogenetic divergence and convergence, emerged at different combinations of phylogenetic locality and abundance weighting. Thus, the ^q^iMPD framework enables phylogenetic scale- and abundance-resolved inference of community assembly. It is implemented in the Python package qdiv version 4.2.0.

## INTRODUCTION

Phylogenetic relationships among community members provide insight into the ecological and evolutionary forces that shape biological communities. Because evolutionary relatedness is often associated with similarity in ecological traits, phylogenetic distances can serve as informative proxies for functional similarity (Martiny et al., 2015; Violle et al., 2011; Washburne et al., 2018). Trait conservation occurs at multiple phylogenetic depths. Complex metabolic processes such as methanogenesis and photosynthesis are conserved in deep lineages, whereas the ability to use simple carbon substrates often varies among shallow, closely related clusters (Martiny et al., 2013). Consequently, the ecological patterns revealed by a metric depend strongly on the phylogenetic scale it emphasizes.

Two metrics, mean pairwise phylogenetic distance (MPD) and mean nearest-taxon distance (MNTD), have become widely used tools for quantifying phylogenetic structure and inferring ecological assembly processes in biological communities (Stegen et al., 2013; Tucker et al., 2017; Webb, 2000; Webb et al., 2002). MPD captures broad evolutionary dispersion by averaging all pairwise distances among taxa, thereby emphasizing deep splits in the phylogeny. MNTD focuses exclusively on each taxon’s closest relative, capturing fine-scale, tip-level patterns. The two metrics also form the basis of the net relatedness index (NRI) and nearest-taxon index (NTI), which express deviations from randomized community assembly as standardized effect sizes relative to null model expectations (Webb et al., 2002). Both metrics have alpha- and beta-diversity formulations, allowing them to quantify phylogenetic dispersion within a community or divergence between communities (Stegen et al., 2012; Swenson, 2011). In microbial ecology, β-MNTD and its standardized effect size, β-NTI, have become widely used for distinguishing deterministic and stochastic assembly processes. By comparing observed phylogenetic turnover to null expectations, β-NTI identifies patterns consistent with homogeneous selection, variable selection, or stochastic community assembly (Dini-Andreote et al., 2015; Stegen et al., 2015).

Despite their widespread use, the existing formulations of the metrics exhibit two major limitations. First, their sensitivity to species abundances is restricted to either presence–absence or linear abundance weighting (Miller et al., 2013; Webb et al., 2008). This is problematic because biological communities typically exhibit highly skewed abundance distributions with a few dominant and many rare taxa; analyses therefore require flexible control over how strongly rare versus abundant taxa should contribute to phylogenetic structure. Second, MPD and MNTD remain conceptually isolated endpoints. MPD reflects global phylogenetic dispersion, while MNTD captures only local, nearest-taxon structure. No existing framework permits continuous tuning between these extremes, nor quantifies the effective phylogenetic neighborhood contributing to the observed pattern.

Here we address both limitations by extending MPD and MNTD into a unified framework. We introduce a locality parameter (*κ*) that adjusts phylogenetic focus from deep to shallow lineages, allowing the classical MPD and MNTD to serve as opposite endpoints of a continuous metric. We adopt the Hill-number weighting scheme, allowing sensitivity to species abundances to be tuned using the diversity order (*q*). The resulting *q*-weighted interpolated mean phylogenetic distance (^q^iMPD) spans a continuum from global to local phylogenetic structure, and from rare to abundant taxa. In combination with null-model analysis, the framework extends NTI- and β-NTI-based assembly inference beyond nearest-taxon structure, enabling tests of whether phylogenetic clustering, overdispersion, convergence, and divergence emerge at different evolutionary depths and abundance weightings. To aid interpretation, we introduce two summary indices, the effective number of neighbors (ENN) and the nearest-taxon focus (NTF), which quantify the effective phylogenetic neighborhood represented by any parameterization. We illustrate the behavior and interpretive value of this framework using an in silico simulated dataset. We then assemble a global dataset of 3661 microbiomes from the wastewater infrastructure and apply the new metric to reveal how phylogenetic structure and community assembly patterns vary among habitats.

## RESULTS

### Abundance weighting modulates phylogenetic structure by shifting emphasis between rare and dominant lineages

To illustrate how the new metric behaves, we used a simulated dataset with 10 species present in three communities (C1-C3). The phylogenetic tree (**Fig. 1a**) includes a shallow clade at the bottom containing species 1-4, and a deep-rooted clade at the top containing species 7-10. The shortest pairwise distance in the tree is 0.15 (between Sp1 and Sp2) and the largest distance is 2.2 (between Sp2 and Sp10). Community composition was designed so that C1 is dominated by species from the shallow clade, C2 is dominated by species from the deep-rooted clade, and C3 contains abundant species from both regions of the tree with Sp1 being the most abundant taxon (**Fig. 1b**).

**Fig 1.**
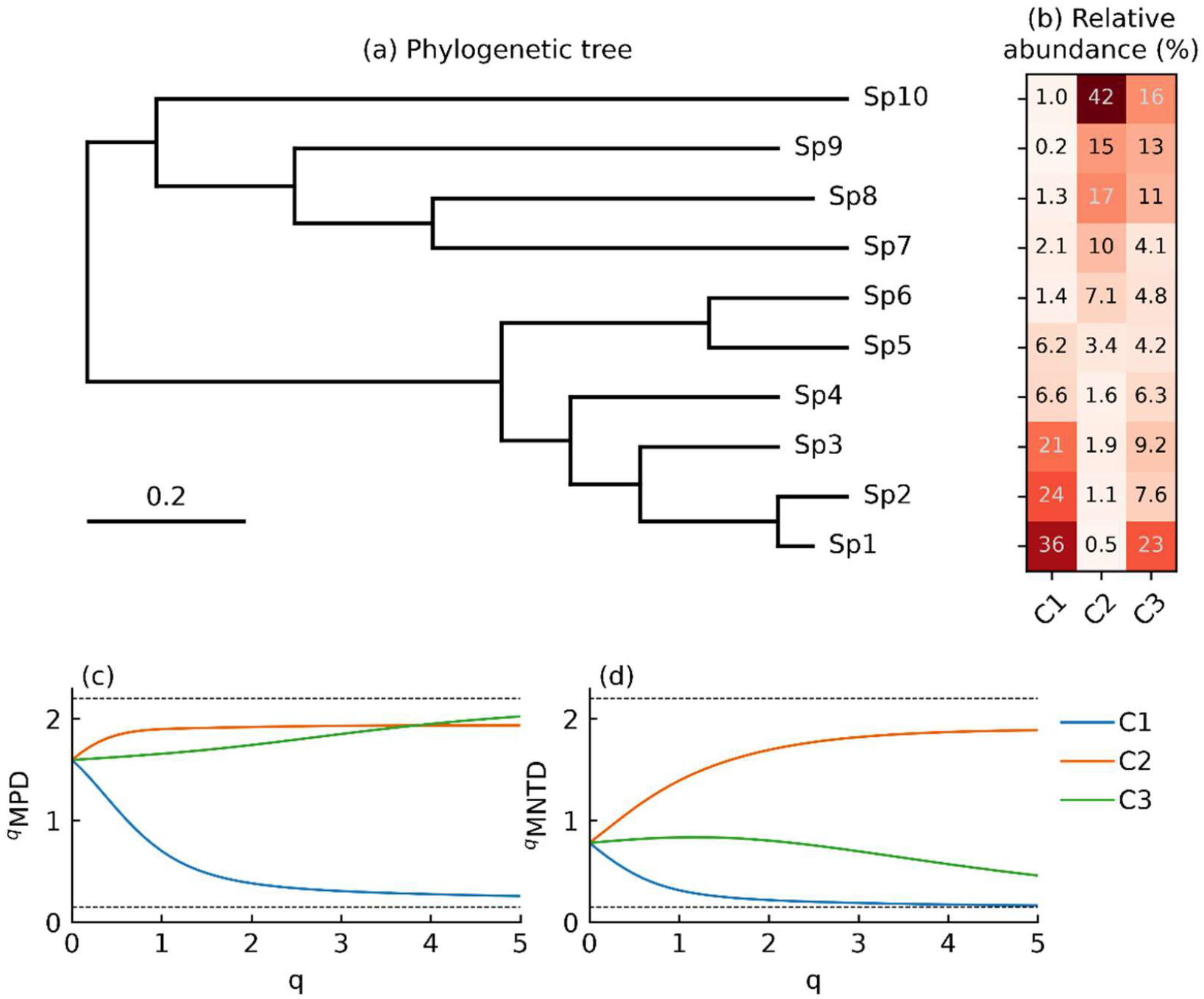
Effect of abundance weighting on mean phylogenetic distance (MPD) and mean nearest-taxon distance (MNTD) in simulated communities. (a) Simulated phylogenetic tree with 10 species, (b) simulated relative abundance distributions of the 10 species in three communities (C1-C3), (c) *q*-weighted MPD, and (d) *q*-weighted MNTD of the three communities.

As the classical MPD and MNTD metrics do not account for species abundances, their values are identical for the three communities. This is shown by the ^0^MPD and ^0^MNTD points in **Fig. 1c–d**. As *q* increases and the metrics give more weight to abundant species, ^q^MPD and ^q^MNTD values for the three communities separate. In C1, both metrics move toward the lower limit of 0.15, which is the distance between the two most abundant species: Sp1 and Sp2. In C2, they approach 1.9, which is the distance between the two most abundant species: Sp9 and Sp10. In C3, ^q^MPD increases with *q*, reflecting the growing influence of Sp1 and Sp10, which occupy opposite ends of the phylogeny and are separated by a distance of 2.15. However, the response of ^q^MNTD differs. It initially increases as the influence of the deeply separated dominant taxa grows but declines again at high *q* as the nearest-neighbor relationships of the most abundant species become increasingly influential. ^q^MPD and ^q^MNTD respond differently to changes in abundance weighting, highlighting their sensitivity to distinct aspects of phylogenetic structure.

### A continuous locality parameter unifies MPD and MNTD and reveals where phylogenetic patterns change

To provide a bridge between MPD and MNTD, we introduced a locality parameter (*κ*) that continuously shifts the phylogenetic scale represented by the ^q^iMPD metric. At *κ*=0, all phylogenetic distances receive equal weight and ^q^iMPD is identical to ^q^MPD. Increasing values of *κ* progressively emphasizes short phylogenetic distances, causing ^q^iMPD to approach the ^q^MNTD limit. **Fig. 2a** shows this transition for combinations of *κ* and *q*-weightings for community C3 in the simulated dataset. At low locality, ^q^iMPD increased with *q*, reflecting the growing influence of the two dominant species, Sp1 and Sp10, which occupy distant regions of the phylogeny. As locality increased, this broad-scale signal weakened because ^q^iMPD became increasingly determined by the nearest phylogenetic neighbors of dominant taxa.

**Fig 2.**
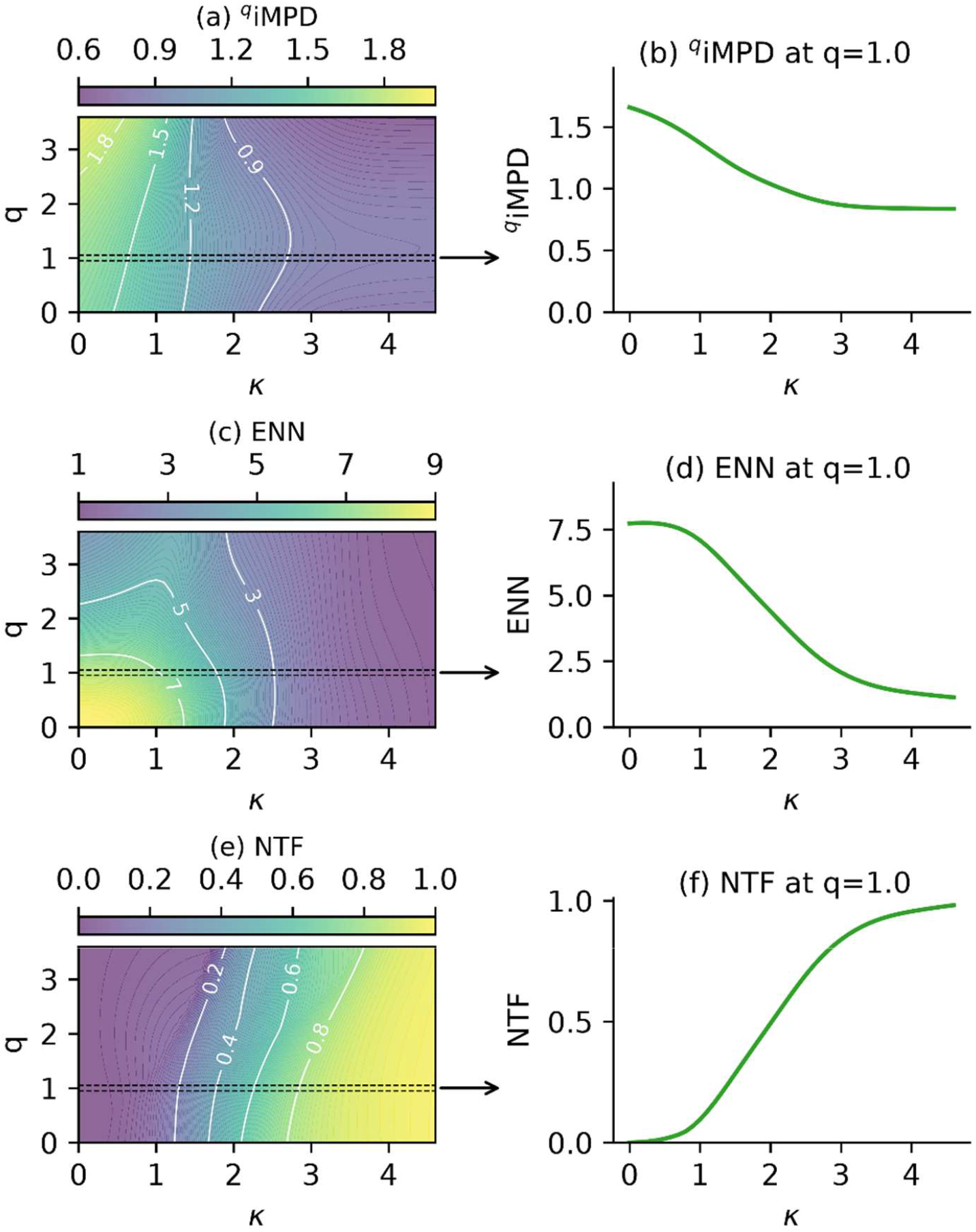
The joint effects of phylogenetic locality (κ) and relative abundance weighting (q) on the interpolated mean phylogenetic distance (^q^iMPD) of community C3 among the simulated communities shown in Fig. 1. (a) ^q^iMPD as functions *κ* and *q*, and (b) as function of *κ* at *q*=1. (c) Effective number of neighbors (ENN) as functions *κ* and *q*, and (d) as function of *κ* at *q*=1. (e) Nearest taxon focus (NTF) as functions *κ* and *q*, and (f) as function of *κ* at *q*=1. The arrows from panels *a, c*, and *e* illustrate that panels *b, d*, and *f* represent a slice of the response surface at *q*=1.

Consequently, the contribution of the large phylogenetic separation between Sp1 and Sp10 was progressively reduced, and the metric became dominated by local phylogenetic structure around the most abundant species.

The slice through the response surface at *q*=1 illustrates how ^q^iMPD varies with locality (**Fig. 2b**). It remained close to its broad-scale value at low *κ*, declined most rapidly at intermediate *κ*, and approached a lower plateau as the metric became dominated by local phylogenetic relationships. Thus, ^q^iMPD does not only combine the MPD and MNTD endpoints, it identifies the range of locality values over which the observed mean phylogenetic distance changes.

To quantify the realized phylogenetic scale underlying ^q^iMPD, we defined the ENN and NTF indices. ENN declined with increasing *κ* showing that fewer neighboring species contributed to the distance calculation as the metric concentrates on short phylogenetic distances (**Fig. 2c-d**). NTF showed the complementary pattern, increasing from zero at *κ* =0 towards one as the calculation approached the nearest-taxon limit (**Fig. 2e-f**). The response surfaces show that the relationship between *κ* and the realized locality depends on abundance weighting (**Fig. 2c, e**). The same value of *κ* can therefore represent different effective phylogenetic neighborhoods at different values of *q*. The behavior of the ^q^iMPD, ENN, and NTF framework for communities C1 and C2 in the simulated dataset is shown in **Fig. S1-S2, supplementary material**.

### Extending the framework to between-community divergence

The ^q^iMPD framework can be extended from within-community dispersion to between-community phylogenetic divergence. **Fig. 3a-c** illustrates the effect of *q*-weighting on β-MPD and β-MNTD for all three pairwise comparisons in the simulated dataset. As *q* increased, the β-^q^MPD metric revealed contrasting patterns among community pairs (**Fig. 3a**). For the C1-C2 comparison, β-^q^MPD increased with *q* as the dominant species occupied opposite ends of the phylogeny. In contrast, β-^q^MPD decreased for C1-C3 because the dominant species were concentrated within the same shallow clade. The C2-C3 comparison showed an intermediate response.

**Fig 3.**
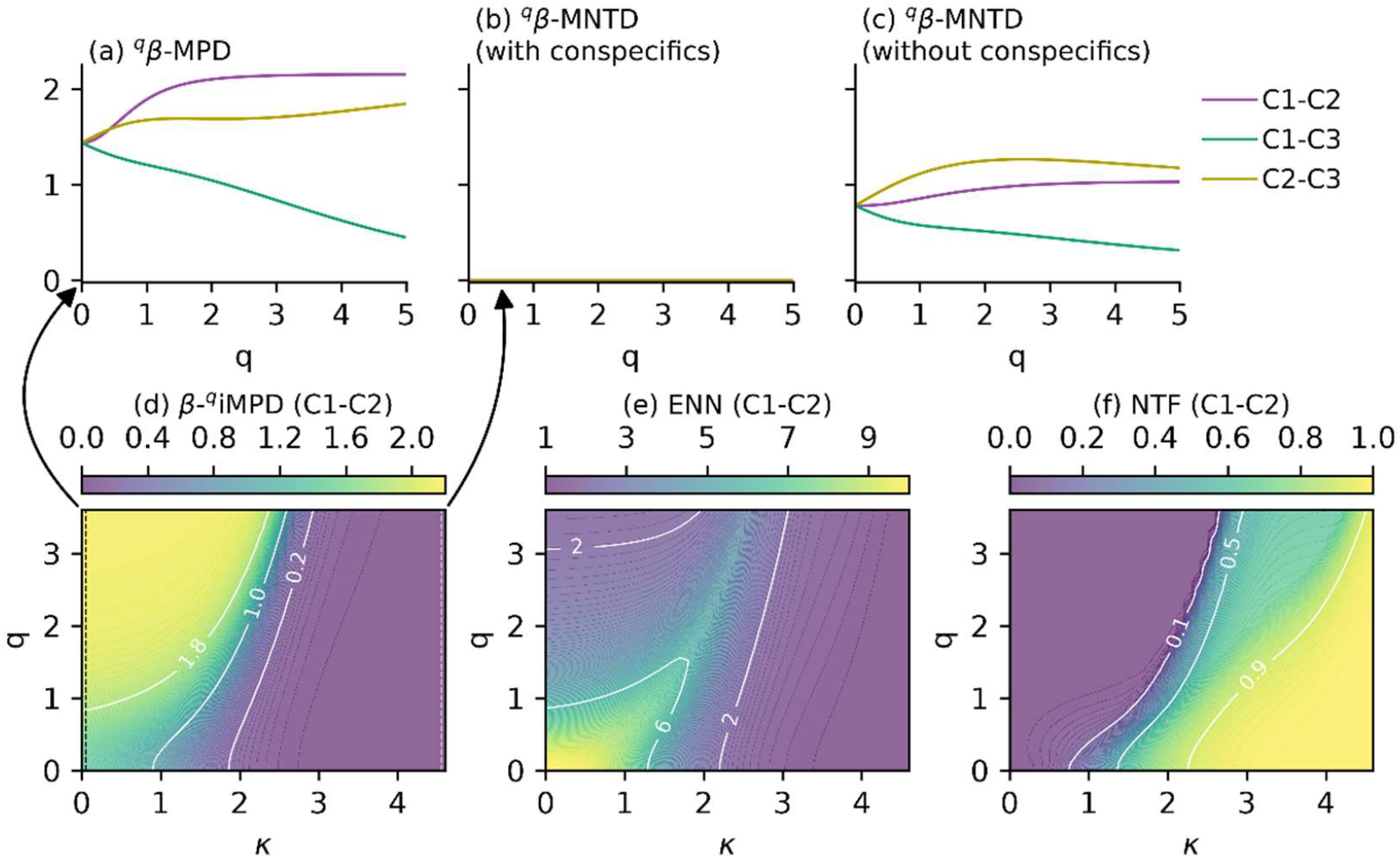
Extension of the framework to divergence between communities. (a) Relative abundance-weighted beta mean phylogenetic distance (β-^q^MPD) and (b-c) beta mean nearest-taxon distance (β-^q^MNTD) for pairwise community comparisons derived from the simulated dataset in Fig. 1. (d) Response surface of β-^q^iMPD for the C1-C2 comparison as a function of relative abundance weighting (*q*) and phylogenetic locality (*κ*), together with the corresponding effective number of neighbors (ENN; e) and nearest taxon focus (NTF; f) response surfaces. The arrows indicate that β-^q^MPD and β-^q^MNTD (with conspecifics included) represent opposite boundaries of a continuous locality spectrum described by β-^q^iMPD.

The behavior of β-^q^MNTD differed substantially. When conspecific matches were included (**Fig. 3b**), β-^q^MNTD collapsed to zero for all pairwise comparisons because all species were present in all three communities. This exposes a fundamental weakness of β-^q^MNTD. When two communities share all species and conspecifics are included in the calculation, β-^q^MNTD necessarily collapses to zero, even when their abundance profiles differ dramatically. Excluding conspecific matches so that the nearest heterospecific taxon is used instead avoids this problem (**Fig. 3c**), but removes an important signal, i.e., the fact that a species is present in both communities should indeed lower their phylogenetic divergence. This limitation is particularly problematic in microbiome studies, where communities contain a vast rare biosphere (Jia et al., 2022; Lynch & Neufeld, 2015). A taxon may be highly abundant in one community and extremely rare in another, yet it contributes the same zero distance to β-^q^MNTD as two equally abundant populations.

The β-^q^iMPD framework provides a more flexible alternative. As locality (*κ*) increases, β-^q^iMPD progressively shifts its emphasis from broad phylogenetic structure toward local nearest-neighbor relationships, while still retaining information from multiple nearby taxa. Consequently, analyses can focus strongly on local phylogenetic structure without requiring a complete collapse to the β-^q^MNTD endpoint. This behavior is illustrated by the response surfaces of the C1-C2 comparison (**Fig. 3d**). At low locality, β-^q^iMPD converged on β-^q^MPD, whereas at high locality it collapsed to zero as the conspecific matches dominated. ENN and NTF showed that the realized locality also depended on *q*, as observed for the within-community metric (**Fig. 3e-f**).

The framework also extends naturally to null-model analyses. Standardized effect sizes can be calculated for both ^q^iMPD and β-^q^iMPD by comparing observed values to null distributions generated by randomizing species identities across the phylogeny while preserving relative abundance structure. This approach generalizes the classical NRI, NTI, and β-NTI frameworks from fixed phylogenetic scales to a continuous spectrum of phylogenetic localities and abundance weightings. In the simulated communities, null model results depended on the selected region of the *κ-q* parameter space (**Fig. S3, supplementary material**). Some communities exhibited consistently clustered or overdispersed phylogenetic structure, whereas others transitioned between clustering and overdispersion as locality and abundance weighting changed. Such transitions were even more common for the pairwise community comparisons, demonstrating that assembly mechanisms inferred from null model analysis can be detected at particular phylogenetic scales and abundance weightings rather than representing fixed properties of a community.

### Phylogenetic structure differs across the wastewater infrastructure

A total of 3661 wastewater microbiomes were categorized as influent, activated sludge, biofilm, aerobic granules, and anaerobic digestion, representing five different habitats in the wastewater treatment infrastructure. For all habitats, abundance weighting had different effects on broad- and local-scale phylogenetic structure (**Fig. 4**). For influent, activated sludge, biofilm, and aerobic granule communities, ^q^MPD was relatively insensitive to abundance weighting, showing changes ranging from a 10.7% decrease to a 0.6% decrease between *q*=0 and *q*=2. In contrast, anaerobic digester communities exhibited a substantial increase in ^q^MPD (16.4%, p<10^-50^). These responses indicate that dominant taxa in digesters are more phylogenetically dispersed than expected from presence-absence information alone, whereas dominant taxa in the other habitats tend to be drawn from more closely related lineages. In contrast to the heterogeneous response of ^q^MPD, ^q^MNTD decreased strongly and consistently across all habitat types.

**Fig 4.**
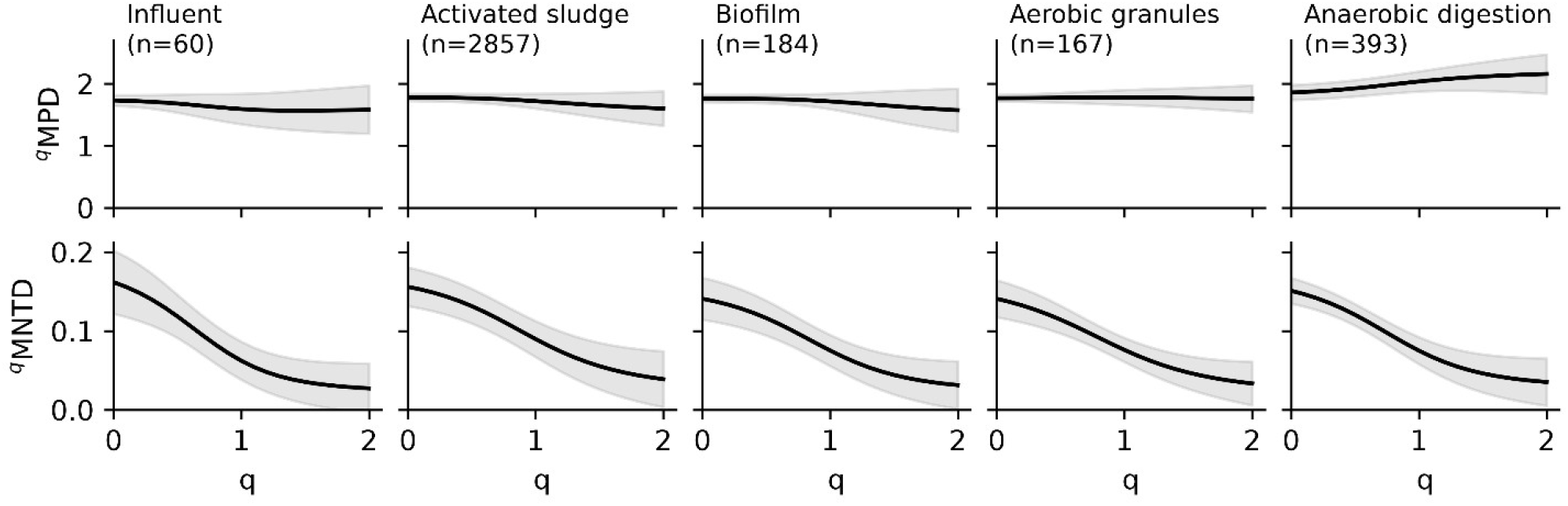
Effect of abundance weighting (*q*) on mean phylogenetic distance (^q^MPD) and mean nearest-taxon distance (^q^MNTD) across five habitats of the wastewater treatment infrastructure. Lines indicate mean values and shaded regions indicate standard deviation among samples. Sample numbers (n) are shown in the panel titles.

Mean ^q^MNTD values were reduced by 75-83% between *q*=0 and *q*=2 (all p<10^-40^). This consistent decline indicates that abundant taxa are generally associated with closer phylogenetic neighbors than rare taxa, regardless of habitat type.

### Community assembly signals depend on both phylogenetic locality and abundance weighting

To evaluate how inferred assembly processes vary with phylogenetic locality and abundance weighting, we applied the framework to two case studies: anaerobic digesters and activated sludge. Null distributions were generated by randomly permuting species identities across the tips of the phylogenetic tree while preserving the observed community abundance structures. The resulting standardized effect sizes (i.e. ^q^iNRI and β-^q^iNRI) quantify deviations from random phylogenetic organization. The within-community metric (^q^iNRI) measures whether taxa in a community are more closely related (phylogenetic clustering) or more distantly related (phylogenetic overdispersion) than expected under the null model. The between-community metric (β-^q^iNRI) evaluates whether pairs of communities are more phylogenetically similar (phylogenetic convergence) or more distinct (phylogenetic divergence) than expected by chance. Positive effect sizes indicate clustering or convergence relative to the null expectation, whereas negative values indicate overdispersion or divergence.

The first case study comprised eight mesophilic and eight thermophilic anaerobic digesters from three countries in the global survey of Dueholm et al. (2024). All digesters were primarily fed with wastewater sludge. The metacommunity, consisting of all species detected in all digesters, included 9148 species. The number of species detected in the individual digester communities ranged from 781 to 1805. At low nearest-taxon focus (NTF), mesophilic digesters exhibited significant phylogenetic overdispersion indicating that coexisting taxa were more distantly related than expected from random assembly (**Fig. 5a**). Thermophilic digesters showed a more complex response, with the inferred assembly pattern depending on abundance weighting. Presence-absence analyses (*q*=0) suggested phylogenetic clustering, whereas abundance-weighted analyses (*q*=1 and *q*=2) indicated weak overdispersion or random assembly (**Fig. 5b**). The broad-scale overdispersion at low NTF is consistent with ecological differentiation among deeply separated phylogenetic lineages, where taxa associated with distinct metabolic functions coexist within anaerobic digesters.

**Fig 5.**
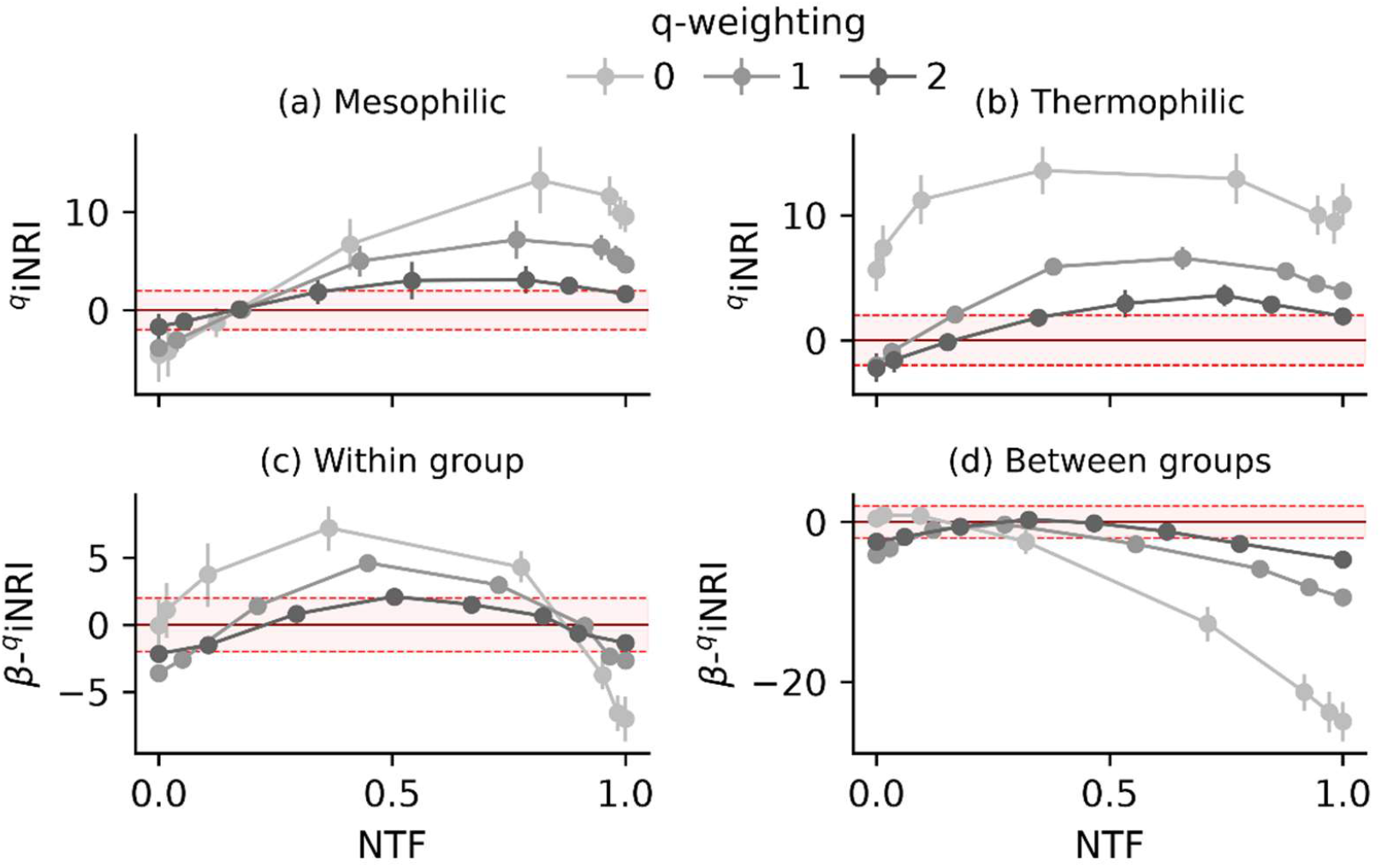
Locality- and abundance-dependent phylogenetic assembly signals in 16 anaerobic digester communities. Standardized effect sizes (*q*-weighted interpolated net relatedness index, ^q^iNRI) were calculated by comparing observed ^q^iMPD and β-^q^iMPD values to null distributions generated by permuting species identities across the phylogeny while preserving community abundance structure. (a-b) Mean and standard deviation of ^q^iNRI values for the mesophilic and thermophilic digester communities. (c) Mean and standard deviation of β-^q^iNRI values for pairwise comparison within the mesophilic and thermophilic groups and (d) between the two groups. Positive values indicate phylogenetic clustering or convergence relative to the null expectation, whereas negative values indicate overdispersion or divergence. Horizontal reference lines mark standardized effect sizes of ±2, which are typically used to indicate statistical significance.

At increasing NTF, both mesophilic and thermophilic communities transitioned towards increasingly positive ^q^iNRI values, indicating a shift from broad-scale phylogenetic overdispersion to local phylogenetic clustering. This transition suggests that different ecological processes dominate at different phylogenetic depths. At broad scales, the coexistence of distantly related taxa is consistent with niche differentiation among major functional groups, whereas at local scales, environmental filtering appears to favor clusters of closely related taxa sharing similar ecological strategies. The strength and position of this transition depended on the abundance-weighting parameter, with presence-absence analyses (*q*=0) generally exhibiting the strongest clustering. These results demonstrate that inferred assembly mechanisms are not fixed properties of anaerobic digester communities but emerge at specific combinations of phylogenetic locality and abundance weighting (**Fig. 5a-b**).

Similar scale-dependent patterns were observed for phylogenetic divergence between communities. Comparisons among digesters of the same temperature regime (mesophilic-mesophilic or thermophilic-thermophilic) yielded positive β-^q^iNRI values at intermediate NTF, indicating greater phylogenetic convergence than expected under the null model. However, this signal weakened and eventually became negative as NTF approached the nearest-neighbor limit, indicating divergence of local phylogenetic neighborhoods (**Fig. 5c**). In contrast, comparisons between mesophilic and thermophilic digesters became progressively more negative with increasing NTF, revealing strong local phylogenetic divergence relative to the null expectation (**Fig. 5d**). These results suggest that temperature has a relatively modest effect on the composition of major phylogenetic lineages but strongly influences the composition of closely related taxa. Consequently, phylogenetic differences between mesophilic and thermophilic digesters are detected most clearly at local phylogenetic scales.

The second case study comprised 20 activated sludge samples collected over a period of two years from a membrane bioreactor studied by de Celis et al. (2022). The metacommunity consisted of 9892 species with between 2336 and 2944 species being detected in each individual sample. To investigate whether temporal succession depended on phylogenetic scale and abundance weighting, we analyzed β-^q^iNRI as a function of sampling interval across the *κ-q* parameter space. The inferred assembly signal varied markedly with phylogenetic locality. At intermediate locality (approximately *κ*=1.5 and NTF=0.5), β-^q^iNRI values were predominantly positive, indicating greater phylogenetic convergence among samples than expected under the null model. In contrast, at high locality (*κ*>2.5, NTF>0.97), β-^q^iNRI values were negative, indicating phylogenetic divergence among the closest phylogenetic neighborhoods (**Fig. 6a**). β-^q^iNRI values were fitted to a model incorporating temporal drift and annual cyclic variation. It was significant only within a restricted subset of parameterizations, with the strongest fits occurring at intermediate to high locality and low to moderate abundance weighting (**Fig. 6b**). The distribution of significant fits showed that the detectability of temporal structure depended on both phylogenetic locality and abundance weighting. Thus, the *κ-q* parameterization at which communities are compared influences not only the inferred assembly pattern but also the ability to detect temporal dynamics (**Fig. 6c-e**).

**Fig 6.**
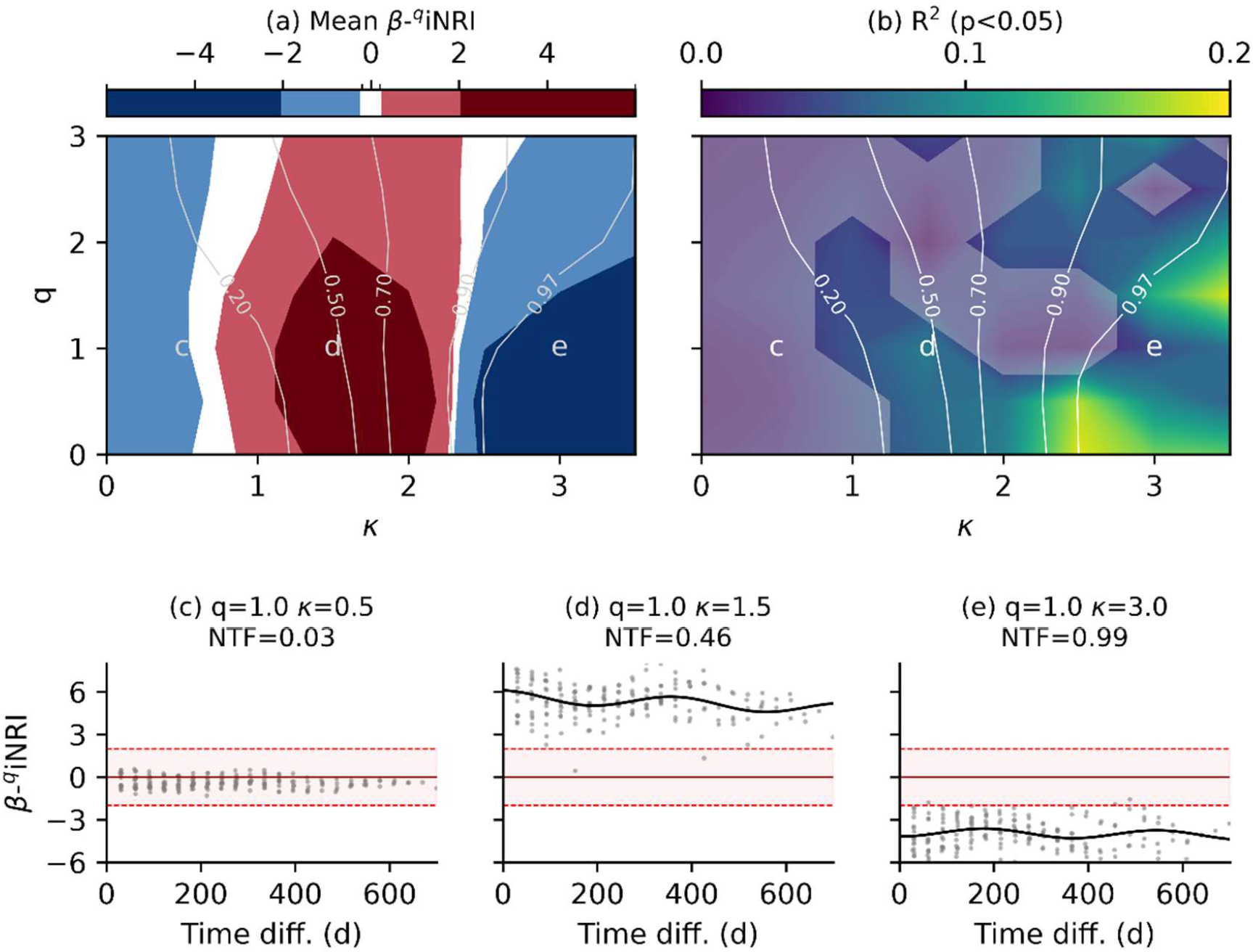
Locality- and abundance-dependent phylogenetic assembly patterns in an activated sludge membrane bioreactor. Standardized effect sizes (*q*-weighted interpolated net relatedness index, β-^q^iNRI) were calculated by comparing observed β-^q^iMPD values with null distributions generated by permuting species identities across the phylogeny while preserving community abundance structure. (a) Mean β-^q^iNRI of all pairwise sample comparisons. (b) Coefficient of determination (R^2^) for model fits. Grey shading indicate parameterizations for which the temporal model fits were not significant (p>0.05). White contour lines in panels a–b indicate mean nearest-taxon focus (NTF). (c-e) β-^q^iNRI as a function of time difference between samples for three *κ–q* parameterizations. Labels c-e in panels a-b indicate the parameterizations shown in panels c-e. (c) Example parameterization with no significant temporal structure at low locality. (d-e) Example parameterizations with significant seasonal variation. Black curves show predictions from multiple regression on matrices (MRM) models containing temporal drift and annual cyclic variation. Red dashed lines indicate standardized effect sizes of ±2.

## DISCUSSION

The ^q^iMPD framework complements existing approaches for incorporating phylogenetic information into diversity analyses. Phylogenetic richness metrics such as Faith’s PD quantify the total amount of evolutionary history represented within communities. Hill-based phylogenetic diversity takes relative abundance information into account to quantify branch diversity or effective number of distinct lineages (Chao et al., 2010). MPD and MNTD, on the other hand, belong to the divergence class of metrics and describe the average phylogenetic separation among taxa (Tucker et al., 2017). With ^q^iMPD, we extend divergence-based phylogenetic analysis into a continuous *q–κ* parameter space, with ENN and NTF describing the realized phylogenetic neighborhood. Researchers can therefore identify where ecological patterns become detectable without selecting a single abundance weighting or choosing categorically between MPD and MNTD.

Applications to wastewater microbiomes showed that abundance weighting and locality expose complementary features of phylogenetic organization. Abundance weighting had relatively little influence on broad-scale dispersion across most habitats but strongly altered local structure, indicating that dominant taxa can be dispersed among major lineages while occupying tightly structured local neighborhoods. The concentration of dominant taxa within tight local neighborhoods may reflect conservation of traits associated with ecological success among closely related taxa, even when those successful groups are distributed across distantly related major lineages. In anaerobic digesters, null-model analyses revealed a clear transition from broad-scale phylogenetic overdispersion to local phylogenetic clustering, suggesting that different assembly processes dominate at different phylogenetic depths. In contrast, between-community analyses showed that inferred patterns of phylogenetic convergence and divergence depended strongly on both locality and abundance weighting. Clustering, overdispersion, convergence, and divergence should therefore not be interpreted as scale-independent properties of communities, but as patterns that emerge at particular combinations of phylogenetic locality and abundance weighting.

The activated sludge time series further showed that the phylogenetic scale of observation can determine both the direction and detectability of assembly patterns. Convergence at intermediate locality gave way to divergence near the nearest-neighbor limit, indicating that temporal turnover was concentrated among closely related taxa rather than among major phylogenetic lineages. However, significant temporal dynamics were detected only within restricted regions of the κ-q space. Phylogenetic locality and abundance weighting can therefore be treated as ecological variables that help identify the evolutionary depths and abundance fractions at which community dynamics and, potentially, the traits underlying them are structured.

An important goal of this work was to make the ^q^iMPD framework accessible. Numba-accelerated implementation in qdiv v4.2.0 makes the framework practical for null-model analyses of species-rich communities such as the wastewater microbiome case studies described above.

## METHODS

### Notation

Let us consider a sample from community *a*. The total number of detected species in the sample is *S*_*a*_ and we denote the set of those species by *A=*⦃a_1_, …, a_S_⦄. The vector of relative abundances is **p**_**a**_ = (p_a1_, …, p_aS_) where is p_ai_ is the relative abundance of species *i* in sample *a*, and 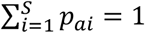. A phylogenetic tree describes the evolutionary relationships among the species. From this tree, we compute a matrix of pairwise phylogenetic distances. We denote this matrix by *D* ∈ ℝ^S×S^, where *S* is the total number of species in the assemblage under study, and each entry in the matrix, *d*_*ij*_, represents the phylogenetic distance between a pair of species (i.e., *i* and *j*). Conspecific distances, i.e., the distance of a species to itself, are zero.

### The classical metrics

The within-community (alpha) variants of MPD and MNTD are shown in Eq. 1a-b. They consider all pairwise phylogenetic distances between species within a community, excluding conspecific distances. The between-community (beta) variants are shown in Eq. 2a-b. They consider all pairwise phylogenetic distances between species detected in one community to species detected in another community, including conspecifics. All four variants of the metrics in Eq. 1-2 can also be abundance weighted by multiplying the distance, *d*_*ij*_, with the relative abundances of species *i* and *j*.

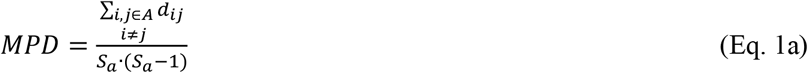

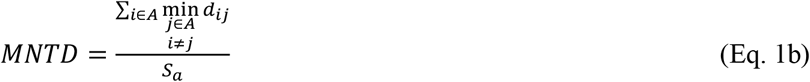

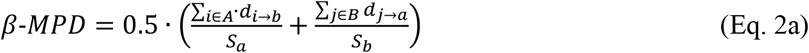

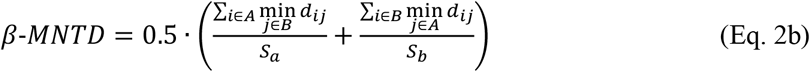

### Relative abundance weighting

To generalize the MPD and MNTD metrics beyond the unweighted and linearly weighted forms, we adopt Hill-number abundance weighting of order *q*. We apply the exponent *q* to the relative abundance vector, **p**, to obtain a normalized *q*-weighted abundance vector, **w** (Eq. 3). If the relative abundance of a species (*p*_*i*_) is zero, the weighted relative abundance (*w*_*i*_) is also zero, irrespective of the value of *q*.

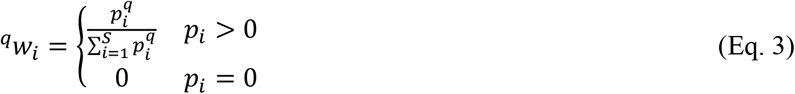

The exponent, *q*, determines the weight given to rare and abundant species. When *q*=0, all present species receive equal weight (^*0*^*w*_*i*_*=1/S*), when *q*=1, the species are weighted exactly according to their relative abundance (^*1*^*w*_*i*_*=p*_*i*_), and larger values of *q* increasingly emphasize more abundant species. For sample *a* with species set *A*, the MPD and MNTD metrics can then be abundance-weighted as shown in Eq. 4a-b.

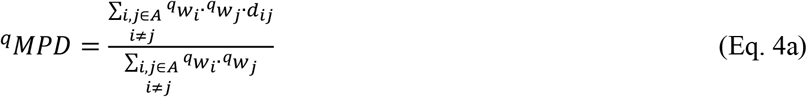

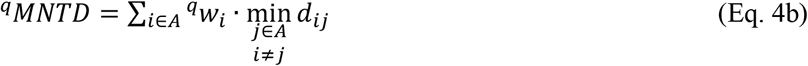

### A continuous bridge between ^q^MPD and ^q^MNTD

To provide a continuous bridge between ^q^MPD (global dispersion) and ^q^MNTD (nearest-neighbor structure), we introduce a soft-minimum kernel with sharpness *r ≥ 0*. The kernel value is calculated for each pairwise distance in *D* (Eq. 5).

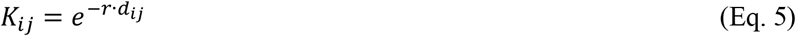

When *r*=0, all distances receive equal weight and the kernel becomes uniform; as *r* increases, the kernel increasingly concentrates on short phylogenetic distances. For a focal species *i*, we convert the kernel into a normalized set of probabilities *Φ* (Eq. 6).

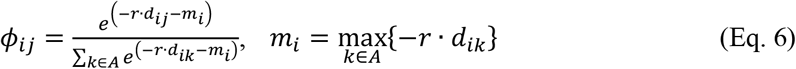

The probability, *Φ*_*ij*_, quantifies how strongly species *i* focuses on the phylogenetic neighborhood of species *j* under a given value of *r*. The kernel is adjusted using the parameter *m*_*i*_ (corresponding to *r* multiplied with the minimum distance in the set) to ensure that the maximum exponent in the kernel is zero, resulting in *K=1*. A soft-minimum distance from species *i* to the rest of the community in sample *a* is defined in Eq. 7, and the ^q^iMPD metric is calculated in Eq. 8.

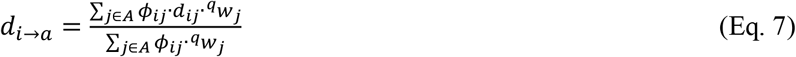

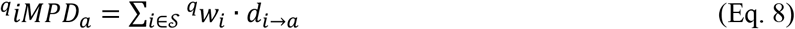

When *r=0*, this formulation yields ^q^*iMPD*_a_ = ^q^*MPD*_a_ and when *r→∞*, it yields 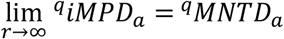.

### A dimensionless locality parameter and two diagnostic indices

As *r* has units of inverse-distance and its effect depend on the magnitude of the values in *D*, we define a scale-free locality parameter, κ, and map it to *r* via the median of the dataset’s distances, excluding conspecifics (Eq. 9).

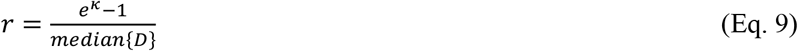

This formulation is designed to give the locality parameter a scale where κ=0 results in uniform kernel values so ^q^iMPD=^q^MPD; κ of approximately 1-2 gives a balanced perspective between ^q^MPD and ^q^MNTD, and higher values of *κ* results in ^q^iMPD approaching ^q^MNTD.

Since *r* is scaled based on the median of distance matrix, the same value of κ may produce different degrees of phylogenetic focus among datasets. To quantify the realized locality of the kernel, we define two diagnostic indices: the effective number of neighbours (ENN) and the nearest taxa focus (NTF). ENN measures the Shannon effective number of neighbors contributing to the soft-minimum distance calculation. It can be interpreted analogously to Hill-number diversity but applied to the neighbor probability distribution rather than species abundances. Values near one indicate that a single neighbor dominates the calculation, whereas larger values indicate contribution from many taxa.

A distribution describing the probability that a neighbor, *j*, contributes to the soft-minimum distance of focal species *i* is described by Eq. 10.

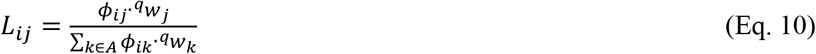

The effective number of neighbors contributing to species *i* is then calculated as the Shannon effective number of the probability distribution (Eq. 11a). Low values of ENN_i_ indicate that only a few neighbors dominate the calculation, whereas large values indicate that many neighbors contribute substantially.

When all probability mass is concentrated on a single neighbor, ENN_i_ = 1. The sample-level effective number of neighbors is calculated according to Eq. 11b.

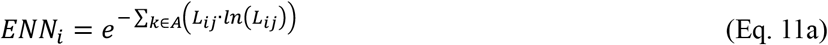

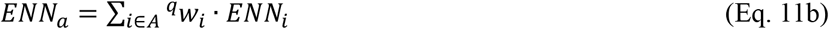

To provide a normalized measure of phylogenetic locality, we define NTF as the position of ENN within its attainable range. The upper endpoint corresponds to a completely diffuse kernel (*r*=0), whereas the lower endpoint corresponds to the nearest-neighbour limit (*r*→∞). NTF therefore varies between 0 and 1, where 0 indicates that the kernel behaves like a global ^q^MPD calculation and 1 indicates that it behaves like a nearest-neighbor ^q^MNTD calculation. Intermediate values represent partial concentration of probability mass on close phylogenetic neighbors.

The maximum value of ENN occurs when locality, *κ*, is zero (Eq. 12a-b).

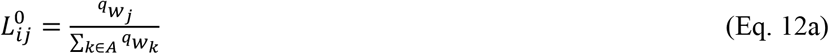

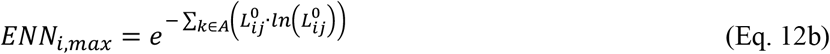

The minimum value of ENN is defined from the set of nearest neighbors (i.e, when *κ* is infinity): *N*_i_ = {*j* ∈ *A* : *d*_ij_ = *d*_i,min_} (Eq. 13a-b).

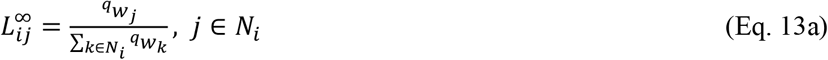

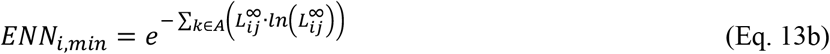

The NTF for species *i* is then calculated by Eq. 14a and the sample-level NTF by Eq. 14b.

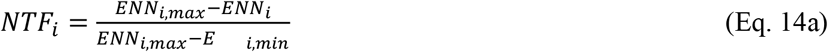

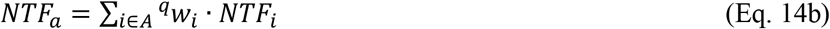

### Extending the interpolated mean phylogenetic distance metric to between-community divergence

For two communities, *a* and *b*, we extend the locality framework to quantify phylogenetic divergence between communities. The directed distance from species *i* in community *a* to the species present in community *b* is calculated using the same kernel formulation as for the within-community case, but with neighboring species weighted by their *q*-weighted relative abundances in community *b* (Eq. 15).

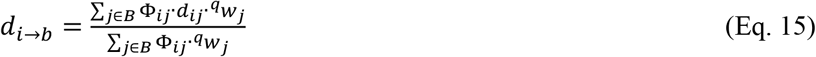

The directed distance, *d*_*i→b*,_ can be interpreted as the kernel-weighted mean phylogenetic distance from species *i* in community *a* to the phylogenetic neighborhood represented by community *b*.

Directed distances are calculated for all species in community *a* and aggregated using the *q*-weighted abundance vector of community *a*. The same procedure is then repeated in the opposite direction, from community *b* to community *a*. The between-community (beta) metric is obtained as the average of the two directed distances (Eq. 16)

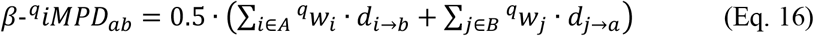

When *κ*=0, the kernel becomes uniform and the metric reduces to β-^q^MPD. As locality increases, progressively greater emphasis is placed on nearby taxa, causing the metric to converge toward the nearest-neighbor limit. Unlike the classical β-^q^MNTD, the β-^q^iMPD framework does not require an arbitrary choice regarding the inclusion or exclusion of conspecific matches. Conspecific taxa contribute naturally to the calculation according to their abundance and the selected locality parameter. At low locality, phylogenetic divergence is determined by the full distribution of phylogenetic distances, whereas at high locality, closely related taxa and conspecific matches receive increasing weight. Consequently, β-^q^iMPD provides a continuous transition between broad-scale and nearest-neighbor perspectives on between-community phylogenetic structure.

### Null model and standardized effect sizes

Just as for the classical MPD and MNTD metrics, null model analysis can also be carried out with the ^q^iMPD metric. We call it *q*-weighted interpolated net relatedness index (^q^iNRI). By permuting species labels across the phylogenetic tree tips or shuffling within-sample abundances, a null model can be generated. The observed ^q^iMPD value can then be compared to the null distribution of values (Eq. 17a). Typically, significance is reported as standardized effect size, where ^q^iNRI > 2 would be considered statistically significant phylogenetic clustering and ^q^iNRI < −2 would be considered statistically significant phylogenetic overdispersion in comparison to the null expectation. The same concept can be used for the between-community metric (Eq. 17b). There, β-^q^iNRI > 2 would be considered statistically significant phylogenetic convergence and β-^q^iNRI < −2 would be statistically significant phylogenetic divergence between communities.

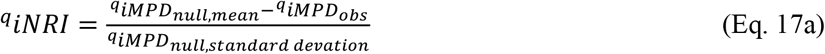

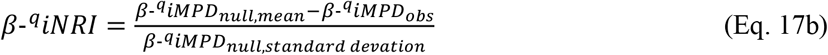

We use the null-minus-observed convention for both ^q^iNRI and β-^q^iNRI so that positive values consistently indicate phylogenetic clustering within communities or convergence between communities. Consequently, it should be noted that the sign convention of β-^q^iNRI is opposite to that commonly used for β-NTI, which is generally calculated as observed β-MNTD minus the mean of the null distribution (Gundersen & Vadstein, 2024; Stegen et al., 2012).

### A global wastewater infrastructure microbiome

For comparability between studies, only studies that had used amplicon sequencing of the V4 hypervariable region of the 16S rRNA gene were included in the study. Sequence reads from 13 studies with samples from 45 countries were downloaded from the sequence read archive (SRA). Primers were removed using cutadapt and paired-end reads were merged using VSEARCH. The trimmed reads were mapped to the Midas Field Guide database v5.3, which contain full-length species representative sequences from the wastewater infrastructure (Nierychlo et al., 2020). Mapping was done with VSEARCH usearch_global using an id threshold of 97% (Rognes et al., 2016). The resulting count table was rarefied to 10 000 reads per sample. The full length sequences from the Midas database were aligned using mafft v7.525 (Katoh et al., 2002), the alignment was trimmed using clipkit v2.13.2 (Steenwyk et al., 2020), and a phylogenetic tree was constructed using FastTree v2.2.0 (Price et al., 2010). Pairwise phylogenetic distances were calculated using qdiv v4.2.0 (Modin, 2026).

Two smaller datasets were used to examine the behavior of ^q^iNRI and β-^q^iNRI in the κ-q parameter space. The anaerobic digestions dataset consisted of 16 communities from the study by Dueholm et al. (2024). The activated sludge dataset consisted of 20 communities from the study by de Celis et al. (2022). In the latter case, triplicate community replicates from each time point were merged prior to analysis. Accession numbers and metadata for all analyzed communities are provided in **File S1 (supplementary material)**.

### Statistical analyses

Changes in ^q^MPD and ^q^MNTD between q=0 and q=2 in the wastewater microbiomes dataset were evaluated by calculating the percentage difference in each metric for every community. Percentage differences within each habitat were then compared against zero using one-sample t-tests implemented in Scipy (Virtanen et al., 2020).

For the activated sludge time series, pairwise β-^q^iNRI values were modeled as a function of temporal drift and annual cyclic variation according to Eq. 18.

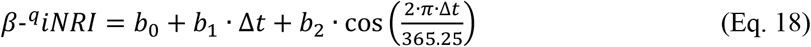

Here, *Δt* is the time difference between two samples (days), and *b*_*0*_, *b*_*1*_, and *b*_*2*_ represent the intercept, temporal drift, and seasonal components, respectively. Model coefficients, R^2^, and significance tests were calculated using the mrm function in qdiv v4.2.0. The function fits the model using ordinary least squares and evaluates statistical significance by permutation analysis (1000 permutations).

### Reproducibility and software

All metrics, diagnostic indices, and null-model procedures are implemented in the Python package qdiv v4.2.0, available from the Python Package Index (PyPI) and at github.com/omvatten/qdiv. Null model calculations use Numba just-in-time compilation, enabling efficient analysis of large sequencing datasets. For the case studies analysed here, comprising approximately 10 000 species and 16–20 communities, 1000-permutation analyses completed in approximately 10–30 min on a standard laptop. Scripts reproducing analyses and figures are provided as **File S2-S5 (supplementary material)**.

