## Supplementary figures for "A unified framework reveals how evolutionary scale and abundance weighting shape phylogenetic community structure"

### Supplementary material

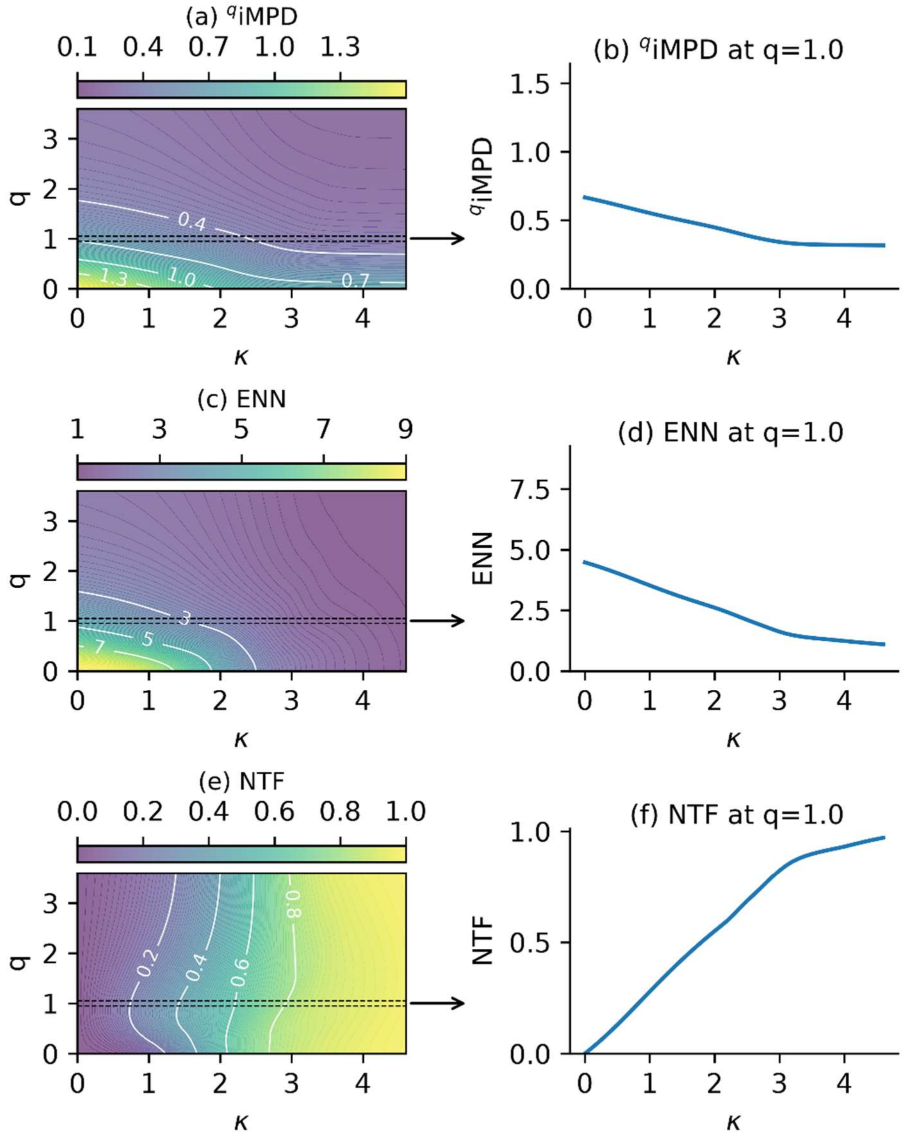


**Fig. S1.** The joint effects of phylogenetic locality (κ) and relative abundance weighting (q) on the interpolated mean phylogenetic distance ($^q$iMPD) of community C1 in the simulated communities shown in Fig. 1. (a) $^q$iMPD as functions κ and q, and (b) as function of κ at q=1. (c) Effective number of neighbors (ENN) as functions κ and q, and (d) as function of κ at q=1. (e) Nearest taxon focus (NTF) as functions κ and q, and (d) as function of κ at q=1. The arrows between from panels *a*, *c*, and *e* illustrate that panels *b*, *d*, and *f* represent a slice of the response surface at q=1.

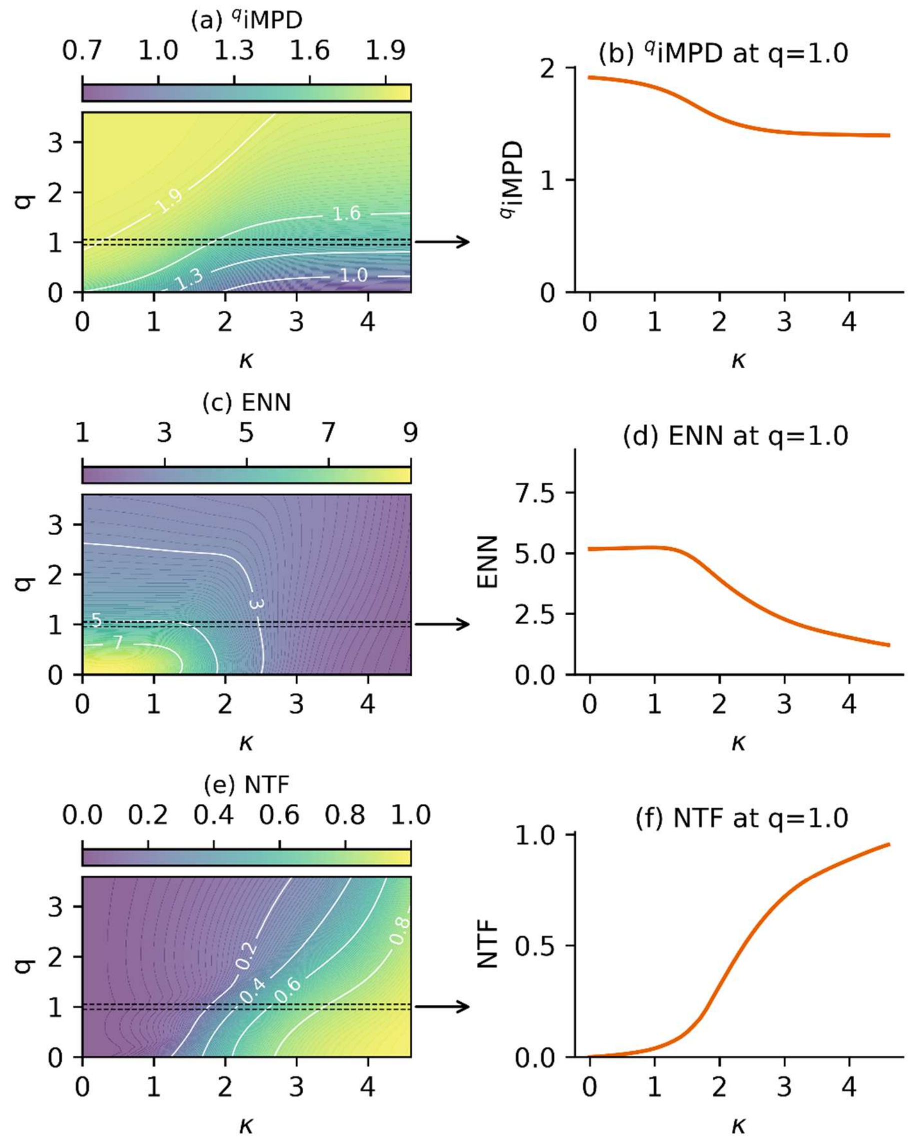


**Fig. S2.** The joint effects of phylogenetic locality (κ) and relative abundance weighting (q) on the interpolated mean phylogenetic distance ($^q$iMPD) of community C2 in the simulated communities shown in Fig. 1. (a) $^q$iMPD as functions κ and q, and (b) as function of κ at q=1. (c) Effective number of neighbors (ENN) as functions κ and q, and (d) as function of κ at q=1. (e) Nearest taxon focus (NTF) as functions κ and q, and (d) as function of κ at q=1. The arrows between from panels *a*, *c*, and *e* illustrate that panels *b*, *d*, and *f* represent a slice of the response surface at q=1.

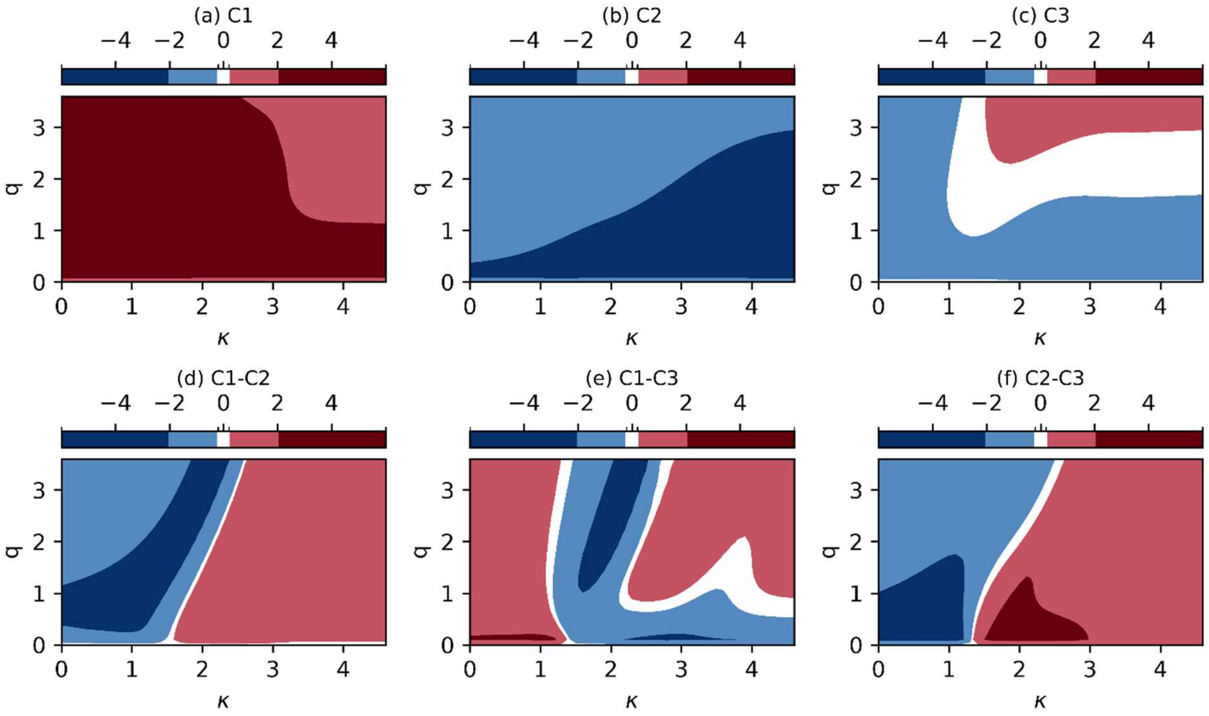


**Fig. S3.** Relative abundance weighted interpolated net relatedness index ($^{q}$iNRI) for the three communities in the simulated dataset (a-c) and β-$^{q}$iNRI for community pairs (d-f). Positive values indicate phylogenetic clustering (a-c) and phylogenetic convergence (d-f) in comparison to random community assembly. Negative values indicate phylogenetic overdispersion or divergence. Values above +2 or below -2 are typically considered statistically significant.
