## Supplementary material for "A unified framework reveals how evolutionary scale and abundance weighting shape phylogenetic community structure": File S2

### Prepare simulated data

#### Create a random imbalanced phylogenetic tree with 10 leaves

In [3]:
```
import qdiv
import numpy as np
import pandas as pd
df, leafs = qdiv.model.make_beta_splitting_tree_df(10, beta=1, random_state=12)
fig, ax, coords = qdiv.plot.phylo_tree({'tree':df, 'leaf_order':leafs},
                                       ladderize=True, invert=False, label_internals=True)
```

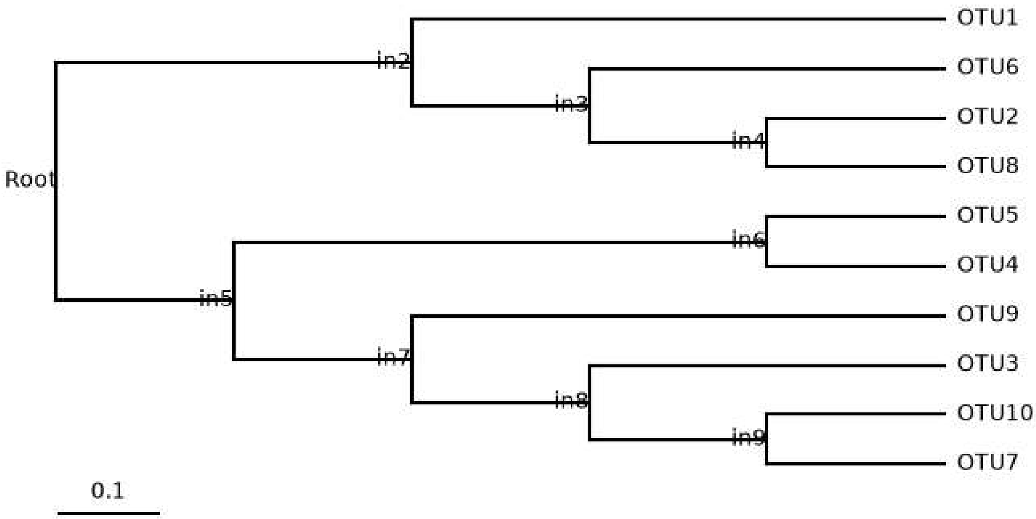


#### Add noise to the branches

In [4]:
```
tip_noise = {
    'in2': -0.3,
    'OTU1': 0.4,
    'OTU8': 0.4,
    'OTU6': 0.3,
    'OTU2': 0.35,
    'in5': 0.4,
    'in6': -0.3,
    'in7': -0.1,
    'in8': -0.1,
    'OTU9': -0.3,
    'OTU3': -0.2,
    'OTU10': -0.1,
    'OTU7': -0.15
}
df, leaves = qdiv.model.make_beta_splitting_tree_df(10, beta=1,
                                                    random_state=12, node_branch_offsets=tip_noise)
fig, ax, coords = qdiv.plot.phylo_tree({'tree':df, 'leaf_order':leaves},
                                       ladderize=True, invert=False, label_internals=True)
```

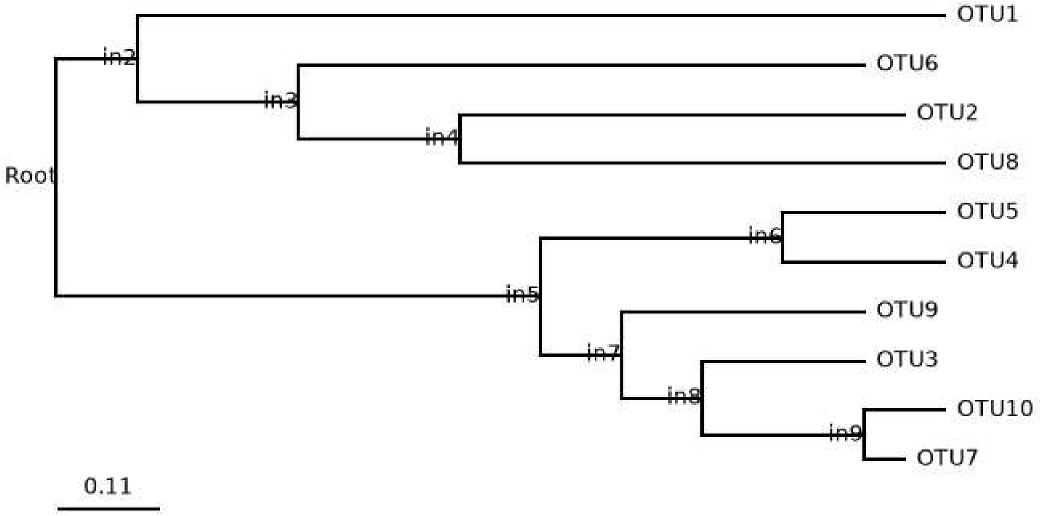


#### Rename leaves, calculate pairwise distance matrix

In [5]:
```
import qdiv
path = 'simulated_data_folder/'
obj = qdiv.MicrobiomeData.load()
obj.tree = df
obj.leaf_order = leaves

obj = obj.rename_features(name_dict={'OTU7': 'Sp1', 'OTU10': 'Sp2',
                                     'OTU3': 'Sp3', 'OTU9': 'Sp4',
                                     'OTU4': 'Sp5', 'OTU5': 'Sp6',
                                     'OTU8': 'Sp7', 'OTU2': 'Sp8',
                                     'OTU6': 'Sp9', 'OTU1': 'Sp10'},
                          inplace=True)
fig, ax, coords = qdiv.plot.phylo_tree(obj, ladderize=True,
                                       invert=False, scale_bar=0.2)

dm = qdiv.sequences.tree_distance_matrix(obj, savename=path+'Simulated_data_distmat')
print('Minimum non-zero distance:\n', dm[dm>0].min())
print('Maximum distance:\n', dm[dm>0].max())
```

```
Tree distance matrix saved to simulated_data_folder\Simulated_data_distmat.csv
Minimum non-zero distance:
 Sp7     1.15
Sp8     1.15
Sp9     1.45
Sp10    1.90
Sp5     0.40
Sp6     0.40
Sp3     0.45
Sp1     0.15
Sp2     0.15
Sp4     0.60
dtype: float64
Maximum distance:
 Sp7     2.20
Sp8     2.15
Sp9     2.10
Sp10    2.20
Sp5     2.20
Sp6     2.20
Sp3     2.10
Sp1     2.15
Sp2     2.20
Sp4     2.10
dtype: float64
```

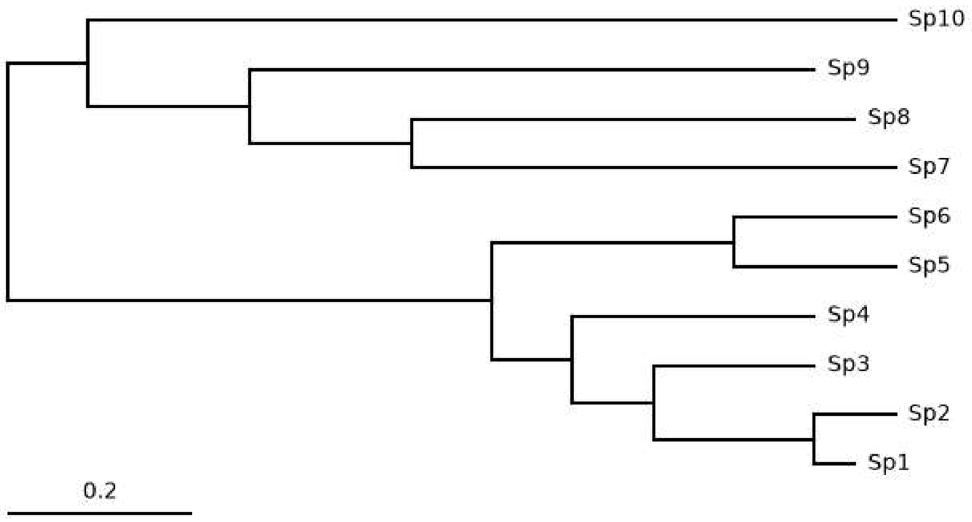


### Generate relative abundance table for three communities

Then save simulated dataset including tree and relative abundance table

In [25]:
```
species = ['Sp'+str(i+1) for i in range(10)]

comm = qdiv.model.simulate_community(size=11, sigma=1.9, random_state=11)
arr1 = qdiv.model.community_sample(comm, n=1000, random_state=20)
arr1 = arr1['Comm1'].to_numpy()[:10]

comm = qdiv.model.simulate_community(size=10, sigma=1.3, random_state=16)
arr2 = qdiv.model.community_sample(comm, n=1000, random_state=21)
arr2 = arr2['Comm1'].to_numpy()[::-1][:10]
print(arr1)
print(arr2)

arr3 = (arr1 + arr2) / 2; arr3[0] -= 20; arr3[-1] += 20

tab = pd.DataFrame({'C1':arr1, 'C2':arr2, 'C3':arr3[::-1]}, index=species)
obj.tab = tab
obj.save(savename=path + 'Simulated')
```

```
[365 240 207  66  62  14  21  13   2  10]
[  5  11  19  16  34  71 105 170 150 419]
```

Out[25]:
```
['simulated_data_folder/Simulated_tab.csv',
 'simulated_data_folder/Simulated_tree.nwk']
```

### Calculate q-profiles for MPD and MNTD

Profiles are saved as pickle file.

In [28]:
```
import qdiv
import numpy as np
import pandas as pd
import pickle
path = 'simulated_data_folder/'
obj = qdiv.MicrobiomeData.load(tab=path+'Simulated_tab.csv',
                               tree=path+'Simulated_tree.nwk')
dm = pd.read_csv(path+'Simulated_data_distmat.csv', index_col=0)

qlist = [round(q, 1) for q in np.arange(0, 5.01, 0.1)]
df = pd.DataFrame(0.0, index=qlist, columns=['C1', 'C2', 'C3'])
df_beta = pd.DataFrame(0.0, index=qlist,
                       columns=['C1-C2', 'C1-C3', 'C2-C3'])
out = {'MPD':df.copy(), 'MNTD':df.copy(),
```

```
        'beta_MPD': df_beta.copy(), 'beta_MNTD': df_beta.copy()
      }

for q in qlist:
    res = qdiv.diversity.mpdq(obj, dm, q=q)
    out['MPD'].loc[q] = res['MPDq']
    res = qdiv.diversity.mntdq(obj, dm, q=q)
    out['MNTD'].loc[q] = res['MNTDq']

    res = qdiv.diversity.beta_mpdq(obj, dm, q=q)
    for s1 in ['C1', 'C2']:
        for s2 in ['C2', 'C3']:
            if s1 != s2:
                s = s1 + '-' + s2
                out['beta_MPD'].loc[q, s] = res.loc[s1, s2]
    res = qdiv.diversity.beta_mntdq(obj, dm, q=q, include_conspecifics=True)
    for s1 in ['C1', 'C2']:
        for s2 in ['C2', 'C3']:
            if s1 != s2:
                s = s1 + '-' + s2
                out['beta_MNTD'].loc[q, s] = res.loc[s1, s2]

with open(path + 'Simulated_MPD_MNTD.pickle', 'wb') as f:
    pickle.dump(out, f)
```

### Plot Figure 1

In [30]:

```
import qdiv
import numpy as np
import pandas as pd
import pickle
import matplotlib.pyplot as plt
import matplotlib.gridspec as gridspec

path = 'simulated_data_folder/'
obj = qdiv.MicrobiomeData.load(tab=path+'Simulated_tab.csv',
                               tree=path+'Simulated_tree.nwk'
                              )
dm = pd.read_csv(path+'Simulated_data_distmat.csv', index_col=0)
dm = dm.loc[obj.tab.index[::-1], obj.tab.index]

with open(path + 'Simulated_MPD_MNTD.pickle', 'rb') as f:
    data = pickle.load(f)

plt.rcParams.update({'font.size':10})
fig = plt.figure(figsize=(20/2.54, 15/2.54))
fig.subplots_adjust(wspace=0.25, hspace=0.5)
gs = gridspec.GridSpec(10, 20)
axT = fig.add_subplot(gs[:6,:15])
axR = fig.add_subplot(gs[:6,16:])
ax1 = fig.add_subplot(gs[7:,0:8])
ax2 = fig.add_subplot(gs[7:,9:17])

#Phylo tree
qdiv.plot.phylo_tree(obj, ax=axT, scale_bar=0.2,
                     scale_bar_y_offset=0.15
                    )

#Relative abundance heatmap
ra = 100 * obj.tab/obj.tab.sum()
im2 = axR.imshow(ra, cmap='Reds', aspect=1)
for y in range(len(ra.index)):
    for x in range(len(ra.columns)):
        val = ra.iloc[y, x]
        if val > 9.9:
            valstr = str(int(round(val, 0)))
        else:
            valstr = str(round(val, 1))
```

```python
        if val > 15:
            col = 'lightgrey'
        else:
            col = 'black'
        axR.text(x, y, valstr, ha='center', va='center',
                 color=col, fontsize=8
                )
axR.set_xticks(range(len(ra.columns)))
axR.set_xticklabels(ra.columns, rotation=45)
axR.set_yticks(range(len(ra.index)))
axR.set_yticklabels([])
axR.invert_yaxis()

cd = {'C1': '#1F78B4', 'C2': '#E66101',
      'C3': '#33A02C'
     }

#MPD
res = data['MPD']
for smp, col in cd.items():
    ax1.plot(res.index, res[smp], color=col,lw=1, label='_nolegend_')

#MNTD
res = data['MNTD']
for smp, col in cd.items():
    ax2.plot(res.index, res[smp], color=col,lw=1, label=smp)

dmin = dm[dm>0].min().min()
dmax = dm.max().max()
for ax in [ax1, ax2]:
    ax.set_xlim(0, 5)
    ax.set_ylim(0, 2.3)
    ax.set_xlabel('q')
    ax.plot(ax.get_xlim(), [dmin, dmin], color='black', ls='--', lw=0.5)
    ax.plot(ax.get_xlim(), [dmax, dmax], color='black', ls='--', lw=0.5)

    for spine in ['top', 'right']:
        ax.spines[spine].set_visible(False)
    for spine in ['left', 'bottom']:
        ax.spines[spine].set_linewidth(0.8)
    ax.tick_params(direction='in', length=3, width=0.8, pad=4)

ax1.set_ylabel('$^q$MPD')
ax2.set_ylabel('$^q$MNTD')
ax1.text(0.1, 2.4, '(c)', ha='left', va='center', fontsize=10)
ax2.text(0.1, 2.4, '(d)', ha='left', va='center', fontsize=10)

ax2.legend(ncol=1, bbox_to_anchor=(1,1), loc=2,
           frameon=False, handletextpad=0.2, columnspacing=1
          )

axT.set_title('(a) Phylogenetic tree', fontsize=10)
axR.set_title('(b) Relative\nabundance (%)', fontsize=10)

plt.savefig(path + 'Figure1_Simulated_qMPD_qMNTD', dpi=600)
plt.savefig(path + 'Figure1_Simulated_qMPD_qMNTD.pdf', format='pdf')
```

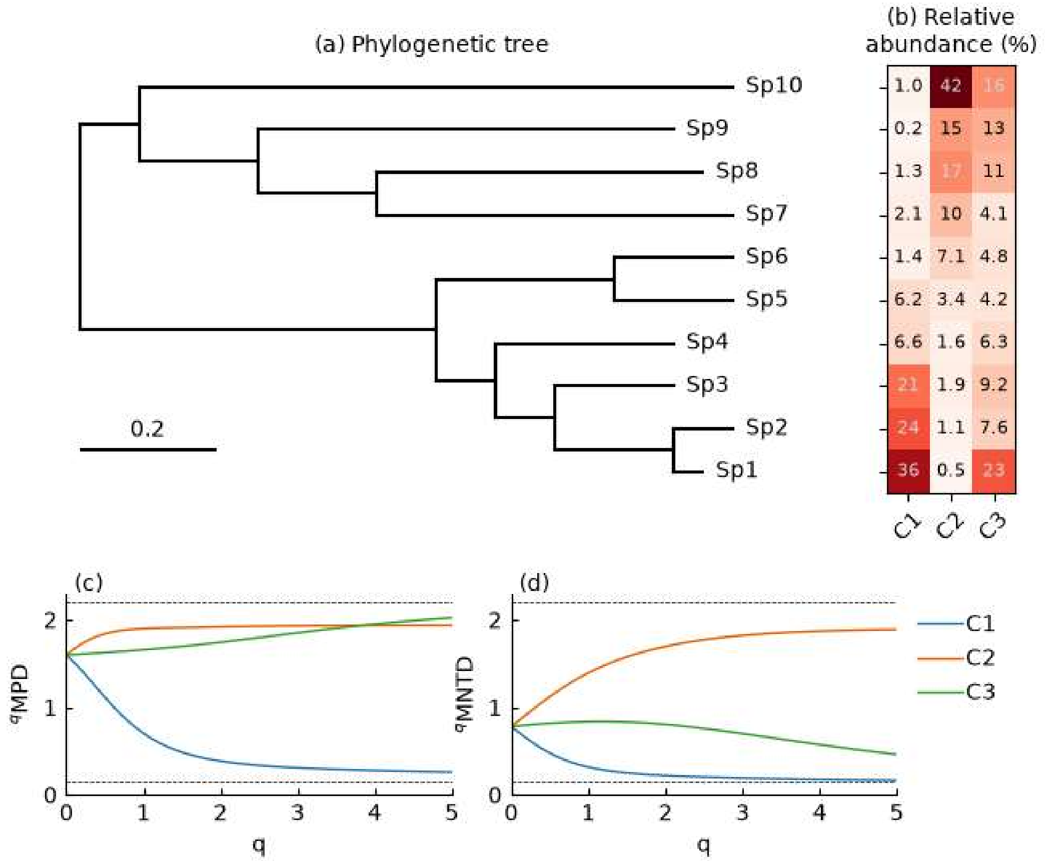


### Calculate locality profiles

In [ ]:

```
import qdiv
import numpy as np
import pandas as pd
import pickle
path = 'simulated_data_folder/'
obj = qdiv.MicrobiomeData.load(tab=path+'Simulated_tab.csv',
                               tree=path+'Simulated_tree.nwk'
                              )
dm = pd.read_csv(path+'Simulated_data_distmat.csv', index_col=0)

qlist = [round(q, 1) for q in np.arange(0, 5.01, 0.1)]
llist = [round(q, 1) for q in np.arange(0, 6.01, 0.1)]
df = pd.DataFrame(0.0, index=llist, columns=['C1', 'C2', 'C3'])

out = {}
for qval in qlist:
    q = round(qval, 1)
    print(q)
    d1 = df.copy()
    d2 = df.copy()
    d3 = df.copy()
    d4 = df.copy()
    for locval in llist:
        loc = round(locval, 1)
        res = qdiv.model.inriq(obj, dm, q=q, locality=loc,
                               iterations=1000, random_state=44
                              )
        d1.loc[loc] = res['iMPDq']
        d2.loc[loc] = res['ENN']
        d3.loc[loc] = res['NTF']
        d4.loc[loc] = res['ses']
    out['iMPD'+str(q)] = d1
```

```
    out['ENN'+str(q)] = d2
    out['NTF'+str(q)] = d3
    out['ses'+str(q)] = d4

with open(path + 'Simulated_iNRIq.pickle', 'wb') as f:
    pickle.dump(out, f)
```

### Figure 2

In [35]:

```
import qdiv
import numpy as np
import pandas as pd
import pickle
import math
import matplotlib.pyplot as plt
import matplotlib.gridspec as gridspec
path = 'simulated_data_folder/'

with open(path + 'Simulated_iNRIq.pickle', 'rb') as f:
    data = pickle.load(f)

q_focus = 1.0
smp_focus = 'C3'
qlim = 3.61; loclim = 4.61
cd = {'C1': '#1F78B4', 'C2': '#E66101', 'C3': '#33A02C'}

qlist = [round(q, 1) for q in np.arange(0, 5.01, 0.1)]
loclist = [round(q, 1) for q in np.arange(0, 6.01, 0.1)]
smplist = ['C1', 'C2', 'C3']
df = pd.DataFrame(0, index=loclist, columns=qlist)
Z = {}; E = {}; N = {}
for s in smplist:
    Z[s] = df.copy()
    E[s] = df.copy()
    N[s] = df.copy()

for q in qlist:
    res1 = data['iMPD'+str(q)]
    res2 = data['ENN'+str(q)]
    res3 = data['NTF'+str(q)]
    for s in smplist:
        Z[s][q] = res1[s]
        E[s][q] = res2[s]
        N[s][q] = res3[s]

plt.rcParams.update({'font.size':10})
fig = plt.figure(figsize=(14/2.54, 17/2.54))
fig.subplots_adjust(wspace=0.5, hspace=0.7)
gs = gridspec.GridSpec(3, 2, width_ratios=[1, 1])
ax1a = fig.add_subplot(gs[0,0])
ax1b = fig.add_subplot(gs[0,1])
ax2a = fig.add_subplot(gs[1,0])
ax2b = fig.add_subplot(gs[1,1])
ax3a = fig.add_subplot(gs[2,0])
ax3b = fig.add_subplot(gs[2,1])

# Fix heatmaps
for a, a2, D, title, title2 in zip(
    [ax1a, ax2a, ax3a],
    [ax1b, ax2b, ax3b],
    [Z, E, N],
    ['(a) $^{q}$iMPD', '(c) ENN', '(e) NTF'],
    ['(b) $^{q}$iMPD at q='+str(q_focus),
     '(d) ENN at q='+str(q_focus),
     '(f) NTF at q='+str(q_focus)
    ]
):
    df = D[smp_focus]
```

```python
    df = df[df.index < loclim]      # κ
    df = df.loc[:, df.columns < qlim]  # q
    X, Y = np.meshgrid(df.index, df.columns)
    Zvals = df.T.to_numpy()
    print(title, Zvals.min(), Zvals.max())
    if 'iMPD' in title:
        zmin = math.floor(Zvals.min()*10) / 10
        zmax = math.ceil(Zvals.max()*10) / 10
        stepsize = (zmax - zmin) / 5
        stepsize = round(stepsize, 1)
        stepslist = [round(x, 1) for x in np.arange(zmin, zmax, stepsize)]
    elif 'ENN' in title:
        zmin = 1; zmax = 9; stepsize = 2
        stepslist = [int(x) for x in np.arange(zmin, zmax+1, stepsize)]
    elif 'NTF' in title:
        zmin = 0; zmax = 1; stepsize = 0.2
        stepslist = [round(x, 1) for x in np.arange(zmin, zmax+0.1, stepsize)]

    cf = a.contourf(X, Y, Zvals, levels=np.linspace(zmin, zmax, 101),
                    cmap='viridis', alpha=0.6
                   )
    cs = a.contour(X, Y, Zvals, levels=np.arange(zmin+stepsize, zmax, stepsize),
                   colors='white', linewidths=0.5
                  )
    a.clabel(cs, fontsize=7) #Contour labels
    a.set_xticks(range(int(loclim)+1))
    a.set_yticks(range(int(qlim)+1))
    a.set_ylabel('q')
    a.set_xlabel(r'$\kappa$')
    cbar = fig.colorbar(cf, ax=a, location='top')
    cbar.set_ticks(stepslist)
    cbar.set_label(title, fontsize=9)

    a.axhline(q_focus-0.05, 0, 1, lw=0.5, ls='--', color='black')
    a.axhline(q_focus+0.05, 0, 1, lw=0.5, ls='--', color='black')
    a.annotate('', xy=(1.2, 1/a.get_ylim()[1]),
               xytext=(1, 1/a.get_ylim()[1]),
               xycoords='axes fraction',
               arrowprops=dict(arrowstyle='->')
              )

    a2.plot(df.index, df[q_focus], color=cd[smp_focus])
    a2.set_ylim(0, Zvals.max()*1.03)
    a2.set_ylabel(title2.split(' ')[1])
    a2.set_xticks(range(int(loclim)+1))
    a2.set_xlabel(r'$\kappa$')
    a2.text(4, a2.get_ylim()[1]*1.02, title2, ha='right')
    for spine in ['top', 'right']:
        a2.spines[spine].set_visible(False)
    for spine in ['left', 'bottom']:
        a2.spines[spine].set_linewidth(0.8)

figname = 'Figure2_simulated_iMPD_q'+str(q_focus).replace('.','')
fig.savefig(path + figname+'_'+smp_focus, dpi=600)
fig.savefig(path + figname+'_'+smp_focus+'.pdf', format='pdf')
```

```
(a) $^{q}$iMPD 0.6224030688899621 1.918729988477151
(c) ENN 1.0505743953158422 9.000000000000002
(e) NTF 0.0 0.9829033596303374
```

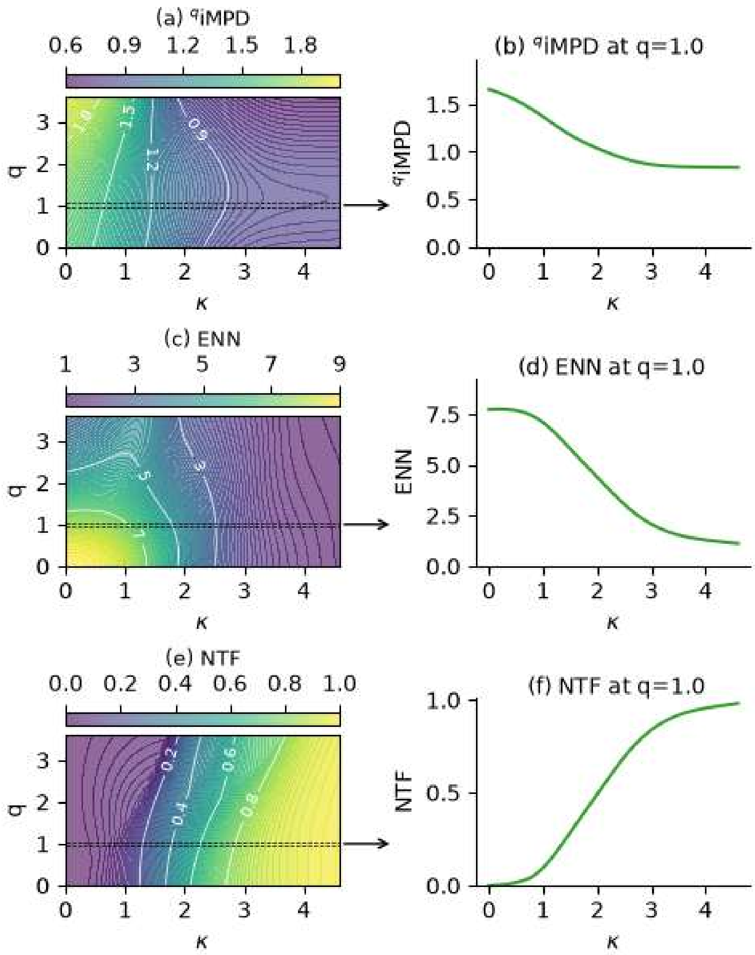


### Calculate beta

In [36]:
```
import qdiv
import numpy as np
import pandas as pd
import pickle

path = 'simulated_data_folder/'
obj = qdiv.MicrobiomeData.load(tab=path+'Simulated_tab.csv',
                               tree=path+'Simulated_tree.nwk'
                               )
dm = pd.read_csv(path+'Simulated_data_distmat.csv', index_col=0)

qlist = [round(q, 1) for q in np.arange(0, 5.01, 0.1)]
smp_pairs = ['C1-C2', 'C1-C3', 'C2-C3']
df_beta = pd.DataFrame(0.0, index=qlist, columns=smp_pairs)
mpd = df_beta.copy()
mntdT = df_beta.copy()
mntdF = df_beta.copy()

for q in qlist:
    res1 = qdiv.diversity.beta_mpdq(obj, dm, q=q)
    res2 = qdiv.diversity.beta_mntdq(obj, dm, q=q, include_conspecifics=True)
    res3 = qdiv.diversity.beta_mntdq(obj, dm, q=q, include_conspecifics=False)
    for sp in smp_pairs:
        s1 = sp.split('-')[0]
        s2 = sp.split('-')[1]
        mpd.loc[q, sp] = res1.loc[s1, s2]
        mntdT.loc[q, sp] = res2.loc[s1, s2]
        mntdF.loc[q, sp] = res3.loc[s1, s2]
```

```
out = {'qMPD':mpd, 'qMNTD_con':mntdT, 'qMNTD_nocon':mntdF}
with open(path + 'Simulated_beta_MPD_MNTD.pickle', 'wb') as f:
    pickle.dump(out, f)
```

In [ ]:

```
import qdiv
import numpy as np
import pandas as pd
import pickle

path = 'simulated_data_folder/'
obj = qdiv.MicrobiomeData.load(tab=path+'Simulated_tab.csv',
                               tree=path+'Simulated_tree.nwk'
                              )
dm = pd.read_csv(path+'Simulated_data_distmat.csv', index_col=0)

qlist = [round(q, 1) for q in np.arange(0, 5.01, 0.1)]
loclist = [round(q, 1) for q in np.arange(0, 6.01, 0.1)]
smp_pairs = ['C1-C2', 'C1-C3', 'C2-C3']
df_beta = pd.DataFrame(0.0, index=loclist, columns=qlist)
df12 = df_beta.copy()
df13 = df_beta.copy()
df23 = df_beta.copy()
n12 = df_beta.copy()
n13 = df_beta.copy()
n23 = df_beta.copy()
e12 = df_beta.copy()
e13 = df_beta.copy()
e23 = df_beta.copy()
i12 = df_beta.copy()
i13 = df_beta.copy()
i23 = df_beta.copy()

for q in qlist:
    for loc in loclist:
        print(q, loc)
        res = qdiv.model.beta_inriq(obj, dm, q=q, locality=loc,
                                    iterations=1000, random_state=44,
                                    use_numba=True
                                   )
        df12.loc[loc, q] = res['beta_iMPDq'].loc['C1', 'C2']
        df13.loc[loc, q] = res['beta_iMPDq'].loc['C1', 'C3']
        df23.loc[loc, q] = res['beta_iMPDq'].loc['C2', 'C3']
        n12.loc[loc, q] = res['beta_NTF'].loc['C1', 'C2']
        n13.loc[loc, q] = res['beta_NTF'].loc['C1', 'C3']
        n23.loc[loc, q] = res['beta_NTF'].loc['C2', 'C3']
        e12.loc[loc, q] = res['beta_ENN'].loc['C1', 'C2']
        e13.loc[loc, q] = res['beta_ENN'].loc['C1', 'C3']
        e23.loc[loc, q] = res['beta_ENN'].loc['C2', 'C3']
        i12.loc[loc, q] = res['beta_ses'].loc['C1', 'C2']
        i13.loc[loc, q] = res['beta_ses'].loc['C1', 'C3']
        i23.loc[loc, q] = res['beta_ses'].loc['C2', 'C3']

out = {'beta12':df12, 'beta13':df13, 'beta23':df23,
       'ntf12':n12, 'ntf13':n13, 'ntf23':n23,
       'enn12':e12, 'enn13':e13, 'enn23':e23,
       'ses12':i12, 'ses13':i13, 'ses23':i23,
       }

with open(path + 'Simulated_beta_iNRIq.pickle', 'wb') as f:
    pickle.dump(out, f)
```

### Figure 3

In [39]:

```
import qdiv
import numpy as np
import pandas as pd
import pickle
```

```python
import matplotlib.pyplot as plt
import matplotlib.gridspec as gridspec
from matplotlib.patches import FancyArrowPatch
path = 'simulated_data_folder/'

cd = {
    'C1': '#1F78B4',
    'C2': '#E66101',
    'C3': '#33A02C',

    'C1-C2': '#984EA3',  # purple
    'C1-C3': '#1B9E77',  # teal
    'C2-C3': '#B2A100'   # olive-gold
}

q_lim = 3.61
loc_lim = 4.61

with open(path + 'Simulated_beta_MPD_MNTD.pickle', 'rb') as f:
    data1 = pickle.load(f)
with open(path + 'Simulated_beta_iNRIq.pickle', 'rb') as f:
    data2 = pickle.load(f)

print(data1.keys())
print(data2.keys())

plt.rcParams.update({'font.size':10})
fig = plt.figure(figsize=(20/2.54, 11/2.54))
fig.subplots_adjust(wspace=1.5, hspace=0.6)
gs = gridspec.GridSpec(2, 34, height_ratios=[1, 1.5])
ax0 = fig.add_subplot(gs[0,:9])
ax1 = fig.add_subplot(gs[0,10:19])
ax2 = fig.add_subplot(gs[0,20:29])

axI = fig.add_subplot(gs[1,:10])
axE = fig.add_subplot(gs[1,12:22])
axN = fig.add_subplot(gs[1,24:34])

for sp in ['C1-C2', 'C1-C3', 'C2-C3']:
    col = cd[sp]
    ax0.plot(data1['qMPD'].index, data1['qMPD'][sp],
             color=col, label=sp, lw=1
            )
    ax1.plot(data1['qMNTD_con'].index, data1['qMNTD_con'][sp],
             color=col, label=sp, lw=1
            )
    ax2.plot(data1['qMNTD_nocon'].index, data1['qMNTD_nocon'][sp],
             color=col, label=sp, lw=1
            )

for ax, let in zip(
    [ax0, ax1, ax2],
    ['(a) $^{q}$$\\beta$-MPD',
     '(b) $^{q}$$\\beta$-MNTD\n(with conspecifics)',
     '(c) $^{q}$$\\beta$-MNTD\n(without conspecifics)'
    ]
):
    ax.set_ylim(-0.02, 2.25)
    ax.set_xlim(0, 5)
    ax.set_xlabel('q')
    ax.text(0.1, 2.3, let, fontsize=9)

    for spine in ['top', 'right']:
        ax.spines[spine].set_visible(False)
    for spine in ['left', 'bottom']:
        ax.spines[spine].set_linewidth(0.8)
    ax.set_xticks(range(6))

ax1.set_yticklabels([])
ax2.set_yticklabels([])
```

```python
ax2.legend(ncol=1, bbox_to_anchor=(1,1), loc=2, frameon=False,
           handletextpad=0.2, columnspacing=1, fontsize=9
          )

Z = 'beta12'; E = 'enn12'; N = 'ntf12';
#Z = 'beta13'; E = 'enn13'; N = 'ntf13';
for a, D, (zmin, zmax), (cmin, cmax, csteps), title in zip(
    [axI, axE, axN], [Z, E, N],
    [(0, 2.2), (0, 10), (0, 1)],
    [(0.2, 2.2, 6), (0, 10, 6), (0.1, 0.9, 3)],
    [r'(d) $\beta$-$^{q}$iMPD (C1-C2)', '(e) ENN (C1-C2)', '(f) NTF (C1-C2)']
):

    df = data2[D]
    df = df[df.index < loc_lim]      # κ
    df = df.loc[:, df.columns < q_lim]  # q
    X, Y = np.meshgrid(df.index, df.columns)
    Zvals = df.T.to_numpy()
    print(Zvals.min(), Zvals.max())

    if 'iMPD' in title:
        zmin = math.floor(Zvals.min()*10) / 10
        zmax = math.ceil(Zvals.max()*10) / 10
        stepsize = (zmax - zmin) / 5
        stepsize = round(stepsize, 1)
        stepslist = [round(x, 1) for x in np.arange(zmin, zmax, stepsize)]
    elif 'ENN' in title:
        zmin = 1; zmax = 10; stepsize = 2
        stepslist = [int(x) for x in np.arange(zmin, zmax+1, stepsize)]
    elif 'NTF' in title:
        zmin = 0; zmax = 1; stepsize = 0.2
        stepslist = [round(x, 1) for x in np.arange(zmin, zmax+0.1, stepsize)]

    cf = a.contourf(X, Y, Zvals, levels=np.linspace(zmin, zmax, 101),
                    cmap='viridis', alpha=0.6
                   )
    cs = a.contour(X, Y, Zvals, levels=np.arange(zmin+stepsize/2, zmax, stepsize*2),
                   colors='white', linewidths=0.5
                  )
    a.clabel(cs, fontsize=7)

    cbar = fig.colorbar(cf, ax=a, location='top')
    cbar.set_ticks(stepslist)
    cbar.set_label(title, fontsize=9)

    a.set_xticks(range(int(loc_lim)+1))
    a.set_xlabel(r'$\kappa$')
    a.set_yticks(range(int(q_lim)+1))
    a.set_ylabel('q')

    if 'MPD' in title:
        bbox1 = a.get_position()
        bbox2 = ax0.get_position()
        bbox3 = ax1.get_position()

        start = (bbox1.x0, bbox1.y0 + bbox1.height)
        end   = (bbox2.x0, bbox2.y0)
        arrow = FancyArrowPatch(
            start, end,
            transform=fig.transFigure,
            connectionstyle="arc3,rad=-0.5",
            arrowstyle="-|>",
            mutation_scale=14,
            lw=1,
            color='black'
        )
        fig.add_artist(arrow)

        start = (bbox1.x1, bbox1.y0 + bbox1.height)
```

```python
        end   = (bbox3.x0+bbox3.width/10, bbox3.y0)
        arrow = FancyArrowPatch(
            start, end,
            transform=fig.transFigure,
            connectionstyle="arc3,rad=0.2",
            arrowstyle="-|>",
            mutation_scale=14,
            lw=1,
            color='black'
        )
        fig.add_artist(arrow)

        xmin, xmax = a.get_xlim()
        a.axvline(xmin+0.05, 0, 1, color='black', ls='--', lw=0.5)
        a.axvline(xmax-0.05, 0, 1, color='lightgrey', ls='--', lw=0.5)

fig.savefig(path + 'Figure3_simulated_beta_iMPD', dpi=600)
fig.savefig(path + 'Figure3_simulated_beta_iMPD.pdf', format='pdf')
```

```
dict_keys(['qMPD', 'qMNTD_con', 'qMNTD_nocon'])
dict_keys(['beta12', 'beta13', 'beta23', 'ntf12', 'ntf13', 'ntf23', 'enn12', 'enn13', 'enn23', 'ses12', 'ses1
3', 'ses23'])
1.8578830549166274e-05 2.1471714010760907
1.0010414600666082 10.000000000000004
0.0 0.9998842822148214
```

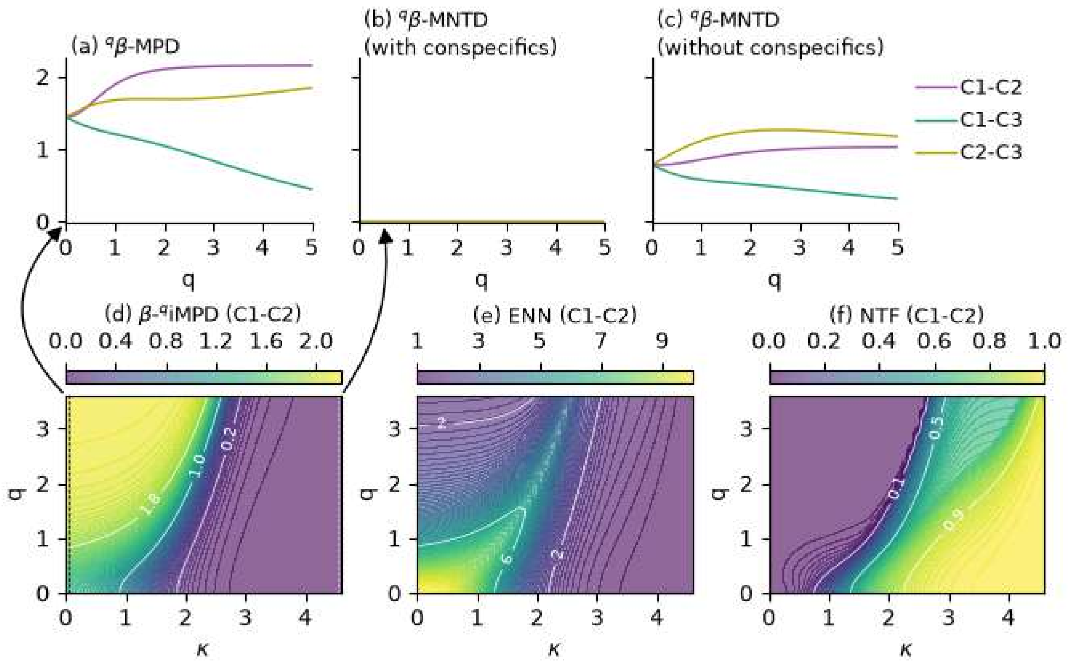


### Plot standardized effect sizes ($^q$iNRI and beta-$^q$iNRI)

In [40]:
```python
import qdiv
import numpy as np
import pandas as pd
import pickle
import matplotlib.pyplot as plt
from matplotlib.colors import LinearSegmentedColormap
from matplotlib.colors import BoundaryNorm

path = 'simulated_data_folder/'

with open(path + 'Simulated_iNRIq.pickle', 'rb') as f:
    data1 = pickle.load(f)
with open(path + 'Simulated_beta_iNRIq.pickle', 'rb') as f:
    data2 = pickle.load(f)
```

```
qlist = [round(q, 1) for q in np.arange(0, 5.01, 0.1)]
loclist = [round(q, 1) for q in np.arange(0, 6.01, 0.1)]

cd = {
    'C1': '#1F78B4',
    'C2': '#E66101',
    'C3': '#33A02C',

    'C1-C2': '#984EA3',  # purple
    'C1-C3': '#1B9E77',  # teal
    'C2-C3': '#B2A100'   # olive-gold
}

df = pd.DataFrame(pd.NA, index=loclist, columns=qlist)
A = {}
for s in ['C1', 'C2', 'C3']:
    A[s] = df.copy()

for q in qlist:
    res = data1['ses'+str(q)]
    for s in res.columns:
        A[s][q] = res[s]

for s, c in zip(['C1-C2', 'C1-C3', 'C2-C3'], ['12', '13', '23']):
    A[s] = data2['ses'+c]

# Plot
plt.rcParams.update({'font.size':10})
fig, axes = plt.subplots(nrows=2, ncols=3, figsize=(20/2.54, 12/2.54))

bounds = [-6, -2, 0, 2, 6]
cmap = LinearSegmentedColormap.from_list(
    'ses',
    [
        '#08306B',  # extreme negative
        '#2166AC',  # -2
        '#FFFFFF',  # -0.5
        '#FFFFFF',  # +0.5
        '#B2182B',  # +2
        '#67000D'   # extreme positive
    ],
    N=256
)
bounds = [-6, -2, -0.2, 0.2, 2, 6]
norm = BoundaryNorm(bounds, cmap.N)

axlist = axes.ravel()
letlist = list('abcdef')
for i, s in enumerate(['C1', 'C2', 'C3',
                       'C1-C2',
                       'C1-C3',
                       'C2-C3'
                      ]
                     ):
    a = axlist[i]
    let = '('+letlist[i]+')'
    df = A[s]
    df = df[df.index < loc_lim]      # κ
    df = df.loc[:, df.columns < q_lim]  # q
    X, Y = np.meshgrid(df.index, df.columns)
    Zvals = df.T.to_numpy()
    print(Zvals.min(), Zvals.max())

    cf = a.contourf(
        X, Y, Zvals,
        levels=np.linspace(-6, 6, 101),
        cmap=cmap,
        norm=norm
    )
```

```
    cbar = fig.colorbar(cf, ax=a, location='top')
    cbar.set_ticks([-4, -2, 0, 2, 4])
    cbar.set_label(let + ' '+s, fontsize=9)

    a.set_xticks(range(int(loc_lim)+1))
    a.set_xlabel(r'$\kappa$')
    a.set_yticks(range(int(q_lim)+1))
    a.set_ylabel('q')

plt.tight_layout()
fig.savefig(path + 'Fig_simulated_ses', dpi=600)
fig.savefig(path + 'Fig_simulated_ses.pdf', format='pdf')
```

```
0.0 3.408224965127313
-2.8306470192330635 0.0
-0.7986875579022327 0.6710786906931356
-2.538423525772037 2.0238857874776146
-2.5564175268388807 2.364095417388986
-3.0293328553063303 2.8200075267582534
```

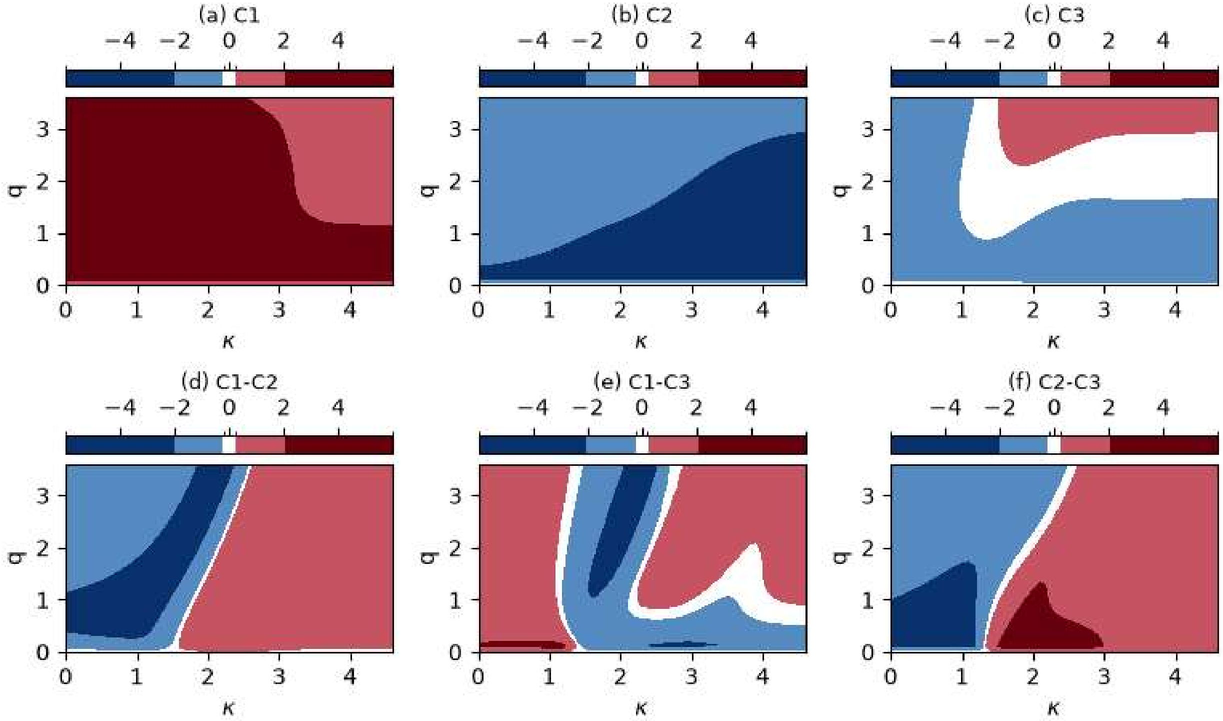


In [ ]:
