## Supplementary material for "A unified framework reveals how evolutionary scale and abundance weighting shape phylogenetic community structure": File S3

### Prepare $^q$MPD and $^q$MNTD for wastewater communities

In [2]:
```
import pandas as pd
import numpy as np
path = 'wastewater_microbiomes_folder/'
meta = pd.read_csv('../16S_data/meta.csv', index_col=0)

meta['cat'] = meta['Process']
meta.loc[meta['Process'].isin(['Biofilter', 'MBBR']), 'cat'] = 'Biofilm'
meta.loc[meta['Process'].isin(['CAS', 'MBR', 'SBR', 'AB_A', 'AB_B']), 'cat'] = 'Activated sludge'
meta.loc[meta['Process'].isin(['Granules', 'AGS']), 'cat'] = 'Aerobic granules'
meta.loc[meta['Process'].isin(['AD']), 'cat'] = 'Anaerobic digestion'

qlist = [round(q, 1) for q in np.arange(0, 2.01, 0.1)]

total = 0
indexlist = []
for c in ['Influent', 'Activated sludge', 'Biofilm', 'Aerobic granules', 'Anaerobic digestion']:
    temp = meta[meta['cat']==c]
    print(c, len(temp))
    total += len(temp)
    indexlist = indexlist + temp.index.tolist()
    df1= pd.DataFrame(pd.NA, index=qlist, columns=temp.index)
    df2= pd.DataFrame(pd.NA, index=qlist, columns=temp.index)
    for s in temp.index:
        res = pd.read_csv('../16S_data/MPD_calc/'+s+'_q.csv', index_col=0)
        df1[s] = res['MPDq']
        df2[s] = res['MNTDq']
    df1.to_csv(path + c.replace(' ','_')+'_MPD.csv')
    df2.to_csv(path + c.replace(' ','_')+'_MNTD.csv')
print('Total number of communities', total)
meta.loc[indexlist].to_csv(path + 'metadata.csv')
```

```
Influent 60
Activated sludge 2857
Biofilm 184
Aerobic granules 167
Anaerobic digestion 393
Total number of communities 3661
```

### Plot figure 4

In [5]:
```
import pandas as pd
import numpy as np
import matplotlib.pyplot as plt
import scipy.stats as stats
path = 'wastewater_microbiomes_folder/'

fut = open(path + 'Fig_MPD_MNTD_all_stats.txt', 'w')

plt.rcParams.update({'font.size':10})
fig, ax = plt.subplots(nrows=2, ncols=5, figsize=(20/2.54, 6.5/2.54),
                       sharex=True, sharey='row',
                       constrained_layout=True
                      )
for i, c in enumerate(['Influent',
                       'Activated sludge',
                       'Biofilm',
                       'Aerobic granules',
                       'Anaerobic digestion'
                      ]
                     ):
    df1 = pd.read_csv(path + c.replace(' ','_')+'_MPD.csv', index_col=0)
    df2 = pd.read_csv(path + c.replace(' ','_')+'_MNTD.csv', index_col=0)
```

```python
    arr1a = df1.iloc[0, :].to_numpy()
    arr1b = df1.iloc[-1, :].to_numpy()
    diff1 = 100 * (arr1b - arr1a) / arr1a
    arr2a = df2.iloc[0, :].to_numpy()
    arr2b = df2.iloc[-1, :].to_numpy()
    diff2 = 100 * (arr2b - arr2a) / arr2a
    res1 = stats.ttest_1samp(diff1, 0)
    res2 = stats.ttest_1samp(diff2, 0)

    fut.write(c + '\n')
    fut.write('MPD % change q2 vs q0 mean='+str(np.mean(diff1))+
              ' std='+str(np.std(diff1))+' pval='+str(res1.pvalue)+'\n')
    fut.write('MNTD % change q2 vs q0 mean='+str(np.mean(diff2))+
              ' std='+str(np.std(diff2))+' pval='+str(res2.pvalue)+'\n')
    fut.write('----------------\n')
    print(c)
    print('MPD percent change q2 vs q0 mean=', round(np.mean(diff1),2))
    print('std=', round(np.std(diff1),2))
    print('pval=', res1.pvalue)
    print('MNTD percent change q2 vs q0 mean=', round(np.mean(diff2),2))
    print('std=', round(np.std(diff2),2))
    print('pval=', res2.pvalue)
    print('------------')

    means1 = df1.mean(axis=1)
    stdevs1 = df1.std(axis=1)

    means2 = df2.mean(axis=1)
    stdevs2 = df2.std(axis=1)

    ax[0, i].fill_between(means1.index,
                          means1-stdevs1, means1+stdevs1,
                          color='grey', alpha=0.2
                         )
    ax[0, i].plot(means1.index, means1, color='black')
    ax[0, i].set_ylim(0, 2.7)
    ax[0, i].text(0.1, 2.5, c + '\n(n='+str(len(df1.columns))+')',
                  fontsize=9
                 )

    ax[1, i].fill_between(means2.index, means2-stdevs2, means2+stdevs2,
                          color='grey', alpha=0.2
                         )
    ax[1, i].plot(means2.index, means2, color='black')

    ax[1, i].set_xlabel('q')
    ax[1, i].set_xticks([0, 1, 2])
    ax[1, i].set_xlim(0, 2.1)

for i in range(2):
    ymin, ymax = ax[i, 0].get_ylim()
    ax[i, 0].set_ylim(0, ymax)
ax[0, 0].set_ylabel('$^q$MPD')
ax[1, 0].set_ylabel('$^q$MNTD')

for a in ax.ravel():
    for spine in ['top', 'right']:
        a.spines[spine].set_visible(False)
    for spine in ['left', 'bottom']:
        a.spines[spine].set_linewidth(0.8)

plt.savefig(path + 'Fig_MPD_MNTD_all_ww', dpi=600)
plt.savefig(path + 'Fig_MPD_MNTD_all_ww.pdf', format='pdf')
fut.close()
```

```
Influent
MPD percent change q2 vs q0 mean= -8.89
std= 21.76
pval= 0.002642432344587999
MNTD percent change q2 vs q0 mean= -83.1
std= 18.66
pval= 1.3172473162411389e-40
------------
Activated sludge
MPD percent change q2 vs q0 mean= -10.24
std= 14.95
pval= 7.024731846241213e-241
MNTD percent change q2 vs q0 mean= -75.39
std= 20.42
pval= 0.0
------------
Biofilm
MPD percent change q2 vs q0 mean= -10.71
std= 19.23
pval= 2.224435908921039e-12
MNTD percent change q2 vs q0 mean= -77.14
std= 22.54
pval= 5.459275722246695e-103
------------
Aerobic granules
MPD percent change q2 vs q0 mean= -0.63
std= 11.11
pval= 0.4628798339279485
MNTD percent change q2 vs q0 mean= -76.79
std= 17.83
pval= 4.337763766418482e-109
------------
Anaerobic digestion
MPD percent change q2 vs q0 mean= 16.4
std= 17.75
pval= 1.7397207113107315e-54
MNTD percent change q2 vs q0 mean= -76.77
std= 19.9
pval= 1.7870495674670228e-237
------------
```

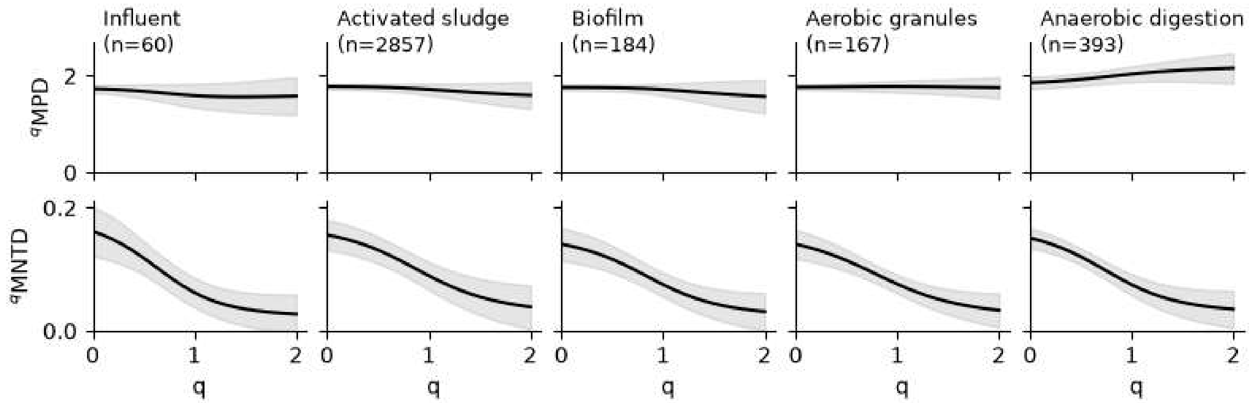
