## Supplementary material for "A unified framework reveals how evolutionary scale and abundance weighting shape phylogenetic community structure": File S4

### Subset data to selected communities

In [ ]:
```
import pandas as pd
import qdiv
path = 'case_AD_folder/'
meta = pd.read_csv(path + 'wastewater_microbiomes_folder/metadata.csv', index_col=0)
m = meta[meta['Case_AD']=='X']
tab = pd.read_csv(path + 'CaseAD_tab.csv')
obj = qdiv.MicrobiomeData.load()
obj.meta = m
obj.tab = tab
obj.add_tree(path + '../16S_data/Detected_tree_rooted.nwk')
obj.prune_tree(featurelist=obj.tab.index.tolist(), inplace=True)
obj.save(path=path, savename='CaseAD'

# Calculate pairwise phylogenetic distances between species
dm = qdiv.sequences.tree_distance_matrix(obj, savename=path + 'CaseAD_distmat', file_format='compressed')
```

### Calculate $^q$iNRI and $\beta$-$^q$iNRI

In [ ]:
```
import qdiv
import pickle
import numpy as np
path = 'case_AD_folder/'

#Load relative abundance table and pre-computed distance matrix
obj = qdiv.MicrobiomeData.load(path=path, tab='CaseAD_tab.csv', meta='CaseAD_meta.csv')
dm = qdiv.sequences.load_compressed_matrix(path + 'CaseAD_distmat.npz')
print('Min', min(dm[dm>0].min())) #Minimum non-zero pairwise distance
print('Max', max(dm[dm>0].max())) #Maximum pairwise distance
print('Median', np.median(dm[dm>0].median())) #Median non-zero pairwise distance

qlist = [0.0, 1.0, 2.0, 3.0, 4.0]
loclist = [0.0, 0.5, 1.0, 1.5, 2.0, 2.5, 3.0]

for q in qlist:
    print('q=', q)
    res = qdiv.model.nriq(obj, distmat=dm, q=q, iterations=1000, random_state=44)
    with open(path + 'NRIq'+str(q)+'.pickle', 'wb') as f:
        pickle.dump(res, f)

    res = qdiv.model.ntiq(obj, distmat=dm, q=q, iterations=1000, random_state=44)
    with open(path + 'NTIq'+str(q)+'.pickle', 'wb') as f:
        pickle.dump(res, f)

    for loc in loclist:
        res = qdiv.model.inriq(obj, distmat=dm, q=q, locality=loc, iterations=1000, random_state=44, use_numba=True)
        with open(path + 'iNRIq'+str(q)+'_loc'+str(loc)+'.pickle', 'wb') as f:
            pickle.dump(res, f)

# Run beta nriq, ntiq, and inriq
for q in qlist:
    print('q=', q)
    res = qdiv.model.beta_nriq(obj, distmat=dm, q=q, iterations=1000, random_state=44)
    with open(path + 'beta_NRIq'+str(q)+'.pickle', 'wb') as f:
        pickle.dump(res, f)

    res = qdiv.model.beta_ntiq(obj, distmat=dm, q=q, iterations=1000, random_state=44, include_conspecifics=True)
    with open(path + 'beta_NTIq'+str(q)+'.pickle', 'wb') as f:
        pickle.dump(res, f)

    for loc in loclist:
        res = qdiv.model.beta_inriq(obj, distmat=dm, q=q, locality=loc, iterations=1000, random_state=44, use_numba=True)
        with open(path + 'beta_iNRIq'+str(q)+'_loc'+str(loc)+'.pickle', 'wb') as f:
            pickle.dump(res, f)
```

### Summarize data for plotting

In [ ]:
```
import pandas as pd
import pickle
import qdiv
path = 'case_AD_folder/'
```

```python
meta = pd.read_csv(path + 'CaseAD_meta.csv', index_col=0)
gr = meta.groupby('Treatment').first()
qlist = [0.0, 1.0, 2.0, 3.0, 4.0]
loclist = [0.0, 0.5, 1.0, 1.5, 2.0, 2.5, 3.0, 9.0]

dfTm = pd.DataFrame(pd.NA, index=loclist, columns=qlist)
dfTs = pd.DataFrame(pd.NA, index=loclist, columns=qlist)
dfMm = pd.DataFrame(pd.NA, index=loclist, columns=qlist)
dfMs = pd.DataFrame(pd.NA, index=loclist, columns=qlist)

dfTENN = pd.DataFrame(pd.NA, index=loclist, columns=qlist)
dfMENN = pd.DataFrame(pd.NA, index=loclist, columns=qlist)
dfTNTF = pd.DataFrame(pd.NA, index=loclist, columns=qlist)
dfMNTF = pd.DataFrame(pd.NA, index=loclist, columns=qlist)

dfWm = pd.DataFrame(pd.NA, index=loclist, columns=qlist)
dfWS1 = pd.DataFrame(pd.NA, index=loclist, columns=qlist)
dfWS2 = pd.DataFrame(pd.NA, index=loclist, columns=qlist)
dfBm = pd.DataFrame(pd.NA, index=loclist, columns=qlist)
dfBS1 = pd.DataFrame(pd.NA, index=loclist, columns=qlist)
dfBS2 = pd.DataFrame(pd.NA, index=loclist, columns=qlist)

dfWENN = pd.DataFrame(pd.NA, index=loclist, columns=qlist)
dfBENN = pd.DataFrame(pd.NA, index=loclist, columns=qlist)
dfWNTF = pd.DataFrame(pd.NA, index=loclist, columns=qlist)
dfBNTF = pd.DataFrame(pd.NA, index=loclist, columns=qlist)

for q in qlist:
    print(q)
    for loc in loclist[:-1]:
        with open(path + 'iNRIq'+str(q)+'_loc'+str(loc)+'.pickle', 'rb') as f:
            res = pickle.load(f)
        meta['alpha'] = res['ses']
        df = meta.groupby("Treatment")["alpha"].agg(["mean", "std"])
        df = df.loc[gr.index]
        dfTm.loc[loc, q] = df.loc['Thermophilic', 'mean']
        dfTs.loc[loc, q] = df.loc['Thermophilic', 'std']
        dfMm.loc[loc, q] = df.loc['Mesophilic', 'mean']
        dfMs.loc[loc, q] = df.loc['Mesophilic', 'std']

        meta['alpha'] = res['ENN']
        df = meta.groupby("Treatment")["alpha"].agg(["mean", "std"])
        df = df.loc[gr.index]
        dfTENN.loc[loc, q] = df.loc['Thermophilic', 'mean']
        dfMENN.loc[loc, q] = df.loc['Mesophilic', 'mean']

        meta['alpha'] = res['NTF']
        df = meta.groupby("Treatment")["alpha"].agg(["mean", "std"])
        df = df.loc[gr.index]
        dfTNTF.loc[loc, q] = df.loc['Thermophilic', 'mean']
        dfMNTF.loc[loc, q] = df.loc['Mesophilic', 'mean']

        # Beta
        with open(path + 'beta_iNRIq'+str(q)+'_loc'+str(loc)+'.pickle', 'rb') as f:
            res = pickle.load(f)
        dis = res['beta_ses']
        bs = qdiv.stats.bootstrap_sample_matrix(dis, meta, by='Treatment',
                                                n_boot=1000, alpha=0.317
                                               )
        dfWm.loc[loc, q] = bs['Treatment']['within']['mean']
        dfWS1.loc[loc, q] = bs['Treatment']['within']['ci'][0]
        dfWS2.loc[loc, q] = bs['Treatment']['within']['ci'][1]
        dfBm.loc[loc, q] = bs['Treatment']['between']['mean']
        dfBS1.loc[loc, q] = bs['Treatment']['between']['ci'][0]
        dfBS2.loc[loc, q] = bs['Treatment']['between']['ci'][1]

        dis = res['beta_ENN']
        bs = qdiv.stats.bootstrap_sample_matrix(dis, meta, by='Treatment',
                                                n_boot=1000, alpha=0.317
                                               )
        dfWENN.loc[loc, q] = bs['Treatment']['within']['mean']
        dfBENN.loc[loc, q] = bs['Treatment']['between']['mean']

        dis = res['beta_NTF']
        bs = qdiv.stats.bootstrap_sample_matrix(dis, meta, by='Treatment',
                                                n_boot=1000, alpha=0.317
                                               )
        dfWNTF.loc[loc, q] = bs['Treatment']['within']['mean']
        dfBNTF.loc[loc, q] = bs['Treatment']['between']['mean']
```

```
    #Add NTIq
    with open(path + 'NTIq'+str(q)+'.pickle', 'rb') as f:
        ntiq = pickle.load(f)
    meta['alpha'] = ntiq['ses']
    df = meta.groupby("Treatment")["alpha"].agg(["mean", "std"])
    df = df.loc[gr.index]
    dfTm.loc[9.0, q] = df.loc['Thermophilic', 'mean']
    dfTs.loc[9.0, q] = df.loc['Thermophilic', 'std']
    dfMm.loc[9.0, q] = df.loc['Mesophilic', 'mean']
    dfMs.loc[9.0, q] = df.loc['Mesophilic', 'std']
    dfTENN.loc[9.0, q] = 1.0
    dfMENN.loc[9.0, q] = 1.0
    dfTNTF.loc[9.0, q] = 1.0
    dfMNTF.loc[9.0, q] = 1.0

    with open(path + 'beta_NTIq'+str(q)+'.pickle', 'rb') as f:
        ntiq = pickle.load(f)
    dis = ntiq['beta_ses']
    bs = qdiv.stats.bootstrap_sample_matrix(dis, meta, by='Treatment',
                                             n_boot=1000, alpha=0.317
                                            )
    dfWm.loc[9.0, q] = bs['Treatment']['within']['mean']
    dfWS1.loc[9.0, q] = bs['Treatment']['within']['ci'][0]
    dfWS2.loc[9.0, q] = bs['Treatment']['within']['ci'][1]
    dfBm.loc[9.0, q] = bs['Treatment']['between']['mean']
    dfBS1.loc[9.0, q] = bs['Treatment']['between']['ci'][0]
    dfBS2.loc[9.0, q] = bs['Treatment']['between']['ci'][1]

    dfWENN.loc[9.0, q] = 1.0
    dfBENN.loc[9.0, q] = 1.0
    dfWNTF.loc[9.0, q] = 1.0
    dfBNTF.loc[9.0, q] = 1.0

out = {}
out['T_mean'] = dfTm
out['T_std'] = dfTs
out['M_mean'] = dfMm
out['M_std'] = dfMs
out['T_ENN'] = dfTENN
out['M_ENN'] = dfMENN
out['T_NTF'] = dfTNTF
out['M_NTF'] = dfMNTF

out['W_mean'] = dfWm
out['W_low'] = dfWS1
out['W_high'] = dfWS2
out['B_mean'] = dfBm
out['B_low'] = dfBS1
out['B_high'] = dfBS2

out['W_ENN'] = dfWENN
out['B_ENN'] = dfBENN
out['W_NTF'] = dfWNTF
out['B_NTF'] = dfBNTF

with open(path + 'Alpha_beta_summary.pickle', 'wb') as f:
    pickle.dump(out, f)
```

### Plot figure 5

In [5]:

```
import pandas as pd
import pickle
import matplotlib.pyplot as plt
path = 'case_AD_folder/'

with open(path + 'Alpha_beta_summary.pickle', 'rb') as f:
    d = pickle.load(f)

colors = ['#bdbdbd', '#969696', '#636363', '#252525']
qlist = [0.0, 1.0, 2.0]
loclist = [0.0, 0.5, 1.0, 1.5, 2.0, 2.5, 3.0]

# iNRI vs loc for different q
plt.rcParams.update({'font.size':10})
fig, ax = plt.subplots(nrows=2, ncols=2, figsize=(14/2.54, 9/2.54), sharex=True)

for i, q in enumerate(qlist):
```

```
    ax[0, 0].errorbar(d['M_NTF'][q], d['M_mean'][q], yerr=d['M_std'][q],
                      color=colors[i], lw=1, marker='o', label=int(q), ms=4
                     )
    ax[0, 1].errorbar(d['T_NTF'][q], d['T_mean'][q], yerr=d['T_std'][q],
                      color=colors[i], lw=1, marker='o', ms=4
                     )
    ax[1, 0].errorbar(d['W_NTF'][q], d['W_mean'][q],
                      yerr=[d['W_mean'][q]-d['W_low'][q], d['W_high'][q]-d['W_mean'][q]],
                      color=colors[i], lw=1, marker='o', ms=4
                     )
    ax[1, 1].errorbar(d['B_NTF'][q], d['B_mean'][q],
                      yerr=[d['B_mean'][q]-d['B_low'][q], d['B_high'][q]-d['B_mean'][q]],
                      color=colors[i], lw=1, marker='o', ms=4
                     )

    ax[0, 0].set_title('(a) Mesophilic', fontsize=9)
    ax[0, 1].set_title('(b) Thermophilic', fontsize=9)
    ax[1, 0].set_title('(c) Within group', fontsize=9)
    ax[1, 1].set_title('(d) Between groups', fontsize=9)

for a in ax.ravel():
    a.set_xlim(-0.05, 1.05)
    a.axhline(-2, 0, 1, color='red', lw=0.5, ls='--')
    a.axhline(0, 0, 1, color='darkred', lw=0.5, ls='-')
    a.axhline(2, 0, 1, color='red', lw=0.5, ls='--')
    a.fill_between(a.get_xlim(), -2, 2, color='red', alpha=0.05)
    a.set_ylabel('$^q$iNRI')
    for spine in ['top', 'right']:
        a.spines[spine].set_visible(False)
    for spine in ['left', 'bottom']:
        a.spines[spine].set_linewidth(0.8)

for a in ax.ravel()[:2]:
    a.set_ylabel('$^q$iNRI')
for a in ax.ravel()[2:]:
    a.set_ylabel(r'$\beta$-$^q$iNRI')
    a.set_xlabel('NTF')

handles, labels = ax[0, 0].get_legend_handles_labels()
fig.legend(handles, labels, ncol=3, title='q-weighting',
           bbox_to_anchor=(0.55,1), loc='upper center',
           frameon=False, handletextpad=0.3, columnspacing=0.8
          )
plt.tight_layout(rect=[0, 0, 1, 0.92])
plt.savefig(path + 'Fig_AD', dpi=600)
plt.savefig(path + 'Fig_AD.pdf', format='pdf')
```

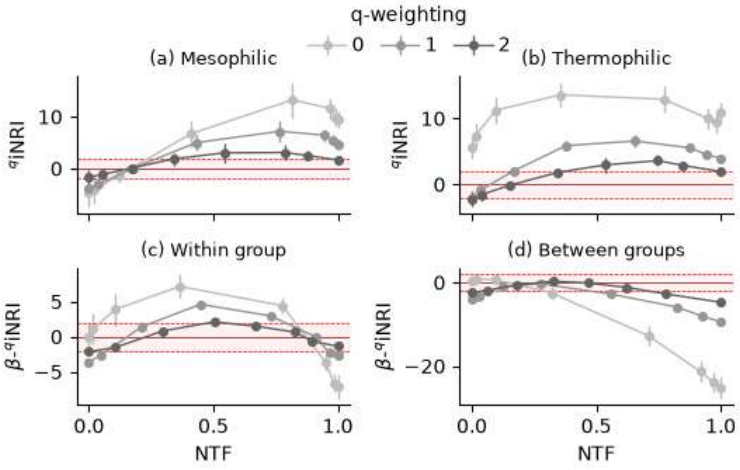
