## Supplementary material for "A unified framework reveals how evolutionary scale and abundance weighting shape phylogenetic community structure": File S5

### Prepare data for MBR case study

In [ ]:
```
import qdiv
path = 'case_MBR_folder/'

obj = qdiv.MicrobiomeData.load(tab=path+'deCelis_tab.csv', meta='wastewater_microbiomes_folder/metadata.csv')

obj.meta['sample_ix'] = obj.meta['Sample_name'].str[:3]
obj = obj.merge_samples(by='sample_ix')
obj = obj.rarefy(depth=30000)

obj.add_tree('../16S_data/Detected_tree_rooted.nwk')
obj.prune_tree(featurelist=obj.tab.index.tolist(), inplace=True)
obj.save(path=path, savename='Case_MBR')
dm = qdiv.sequences.tree_distance_matrix(obj, savename=path + 'CaseMBR_distmat', file_format='compressed')
```

### Calculate ${}^q$iNRI and $\beta$-${}^q$iNRI

In [ ]:
```
import qdiv
import pickle
import numpy as np
path = 'case_MBR_folder/'

obj = qdiv.MicrobiomeData.load(path=path, tab='CaseMBR_tab.csv',
                               meta='CaseMBR_meta.csv'
                              )
dm = qdiv.sequences.load_compressed_matrix(path + 'CaseMBR_distmat.npz')

qlist = [0.0, 0.5, 1.0, 1.5, 2.0, 2.5, 3.0]
loclist = [0.0, 0.5, 1.0, 1.5, 2.0, 2.5, 3.0, 3.5]

for q in qlist:
    print('q=', q)
    for loc in loclist:
        res = qdiv.model.inriq(obj, distmat=dm, q=q, locality=loc,
                               iterations=1000, random_state=44, use_numba=True
                              )
        with open(path + 'iNRIq'+str(q)+'_loc'+str(loc)+'.pickle', 'wb') as f:
            pickle.dump(res, f)

for q in qlist:
    print('q=', q)
    for loc in loclist:
        res = qdiv.model.beta_inriq(obj, distmat=dm, q=q, locality=loc,
                                    iterations=1000, random_state=44, use_numba=True
                                   )
        with open(path + 'beta_iNRIq'+str(q)+'_loc'+str(loc)+'.pickle', 'wb') as f:
            pickle.dump(res, f)
```

### Get an overview of results

In [12]:
```
import pandas as pd
import pickle
import numpy as np
import math
import matplotlib.pyplot as plt
import matplotlib.dates as mdates
import qdiv
path = 'case_MBR_folder/'

saved = {}

meta = pd.read_csv(path + 'CaseMBR_meta.csv', index_col=0)
meta['month'] = meta.index.str[1:].astype(float)

meta['Collection_date'] = pd.to_datetime(meta['Collection_date'])
td = qdiv.stats.pairwise_difference(meta, by='Collection_date')
td = td['Collection_date']
x = td.to_numpy()[np.triu_indices(len(td), k=1)]

start_date = pd.Timestamp("2017-04-01")
meta['day'] = (meta["Collection_date"] - start_date).dt.days
```

```python
timediff = qdiv.stats.pairwise_difference(meta, by='day')['day']
cyc = np.cos(2*math.pi * timediff.to_numpy() / 365.25)
cyc = pd.DataFrame(cyc, index=timediff.index, columns=timediff.columns)

qlist = [0.0, 0.5, 1.0, 1.5, 2.0, 2.5, 3.0]
loclist = [0.0, 0.5, 1.0, 1.5, 2.0, 2.5, 3.0, 3.5]

plt.rcParams.update({'font.size':9})
fig, ax = plt.subplots(nrows=len(loclist), ncols=len(qlist), figsize=(30/2.54, 40/2.54), sharex=True, sharey='col')

for c, q in enumerate(qlist):
    for r, loc in enumerate(loclist):
        print(q, loc)
        sn = 'q='+str(q)+'_loc='+str(loc)
        saved[sn] = {}
        saved[sn]['q'] = q
        saved[sn]['loc'] = loc

        with open(path + 'beta_iNRIq'+str(q)+'_loc'+str(loc)+'.pickle', 'rb') as f:
            res = pickle.load(f)

        dis = res['beta_ses'].loc[td.index, td.columns]
        disv = dis.to_numpy()[np.triu_indices(len(dis), k=1)]
        saved[sn]['ses'] = disv
        saved[sn]['x'] = x

        ntf = res['beta_NTF'].loc[td.index, td.columns]
        ntfv = ntf.to_numpy()[np.triu_indices(len(ntf), k=1)]
        saved[sn]['NTF'] = ntfv

        ax[r, c].scatter(x, disv, marker='.', color='black', s=2)

        st = qdiv.stats.mrm(dis, {'cyc':cyc, 'timediff':timediff})
        for key, item in st.items():
            if isinstance(item, dict):
                print(key, ':')
                for key2, item2 in item.items():
                    print(key2, '=')
                    print(item2)
            else:
                print(key, '=')
                print(item)
        print('---')
        saved[sn]['R2'] = st['R2']
        saved[sn]['R2_p'] = st['R2_p']
        saved[sn]['cyc_p'] = st['coefficient_pvals']['cyc']
        saved[sn]['timediff_p'] = st['coefficient_pvals']['timediff']
        saved[sn]['cyc'] = st['coefficients']['cyc']
        saved[sn]['timediff'] = st['coefficients']['timediff']

        title = r'q='+str(q)+' $\kappa$='+str(loc)
        title = title +' NTF='+str(round(np.mean(ntfv),2))
        title = title + '\nR2='+str(round(st['R2'],2))

        if st['R2_p'] < 0.05:
            ax[r, c].set_title(title+'*', fontsize=8)
            xrange = np.linspace(0, 700, 200)
            yrange = np.cos(2*math.pi * xrange / 365.25) * st['coefficients']['cyc']
            yrange = yrange + xrange * st['coefficients']['timediff']
            yrange = yrange + st['coefficients']['Intercept']
            ax[r, c].plot(xrange, yrange, color='magenta', lw=1)
            saved[sn]['xrange'] = xrange
            saved[sn]['yrange'] = yrange
        else:
            ax[r, c].set_title(title, fontsize=8)

        ax[r, c].axhline(-2, 0, 1, color='red', lw=0.5, ls='--')
        ax[r, c].axhline(0, 0, 1, color='darkred', lw=0.5, ls='-')
        ax[r, c].axhline(2, 0, 1, color='red', lw=0.5, ls='--')
        ax[r, c].fill_between([min(x), max(x)], -2, 2, color='red', alpha=0.05)

for i in range(len(qlist)):
    ax[-1, i].set_xlabel('Time diff. (d)')
plt.tight_layout()
plt.savefig(path + 'Fig_beta_iNRIq_vs_sampling_time', dpi=600)

with open(path + 'Stats_beta.pickle', 'wb') as f:
    pickle.dump(saved, f)
```

```
0.0 0.0
coefficients :
Intercept =
-0.04314111709086586
cyc =
0.13718153089980412
timediff =
-0.001229063659399651
coefficient_pvals :
Intercept =
0.94
cyc =
0.146
timediff =
0.376
R2 =
0.015329894077701911
R2_p =
0.324
---
0.0 0.5
coefficients :
Intercept =
0.07210758891260674
cyc =
0.2105250693475868
timediff =
-0.001637759400590987
coefficient_pvals :
Intercept =
0.898
cyc =
0.049
timediff =
0.294
R2 =
0.02306203557141151
R2_p =
0.22
---
0.0 1.0
coefficients :
Intercept =
1.1601947454943868
cyc =
0.3971979565307539
timediff =
-0.0026439817828363475
coefficient_pvals :
Intercept =
0.081
cyc =
0.005
timediff =
0.176
R2 =
0.044325192430156646
R2_p =
0.091
---
0.0 1.5
coefficients :
Intercept =
4.210688967924597
cyc =
0.8270657919732611
timediff =
-0.004471848990021817
coefficient_pvals :
Intercept =
0.046
cyc =
0.001
timediff =
0.111
R2 =
0.06654659685703823
R2_p =
0.036
---
```

```
0.0 2.0
coefficients :
Intercept =
4.436129107408658
cyc =
1.1930207180129073
timediff =
-0.004380286867471271
coefficient_pvals :
Intercept =
0.066
cyc =
0.001
timediff =
0.121
R2 =
0.08585057363218829
R2_p =
0.009
---
0.0 2.5
coefficients :
Intercept =
-1.4722541135442346
cyc =
1.079047195987157
timediff =
-0.004195279331045367
coefficient_pvals :
Intercept =
0.997
cyc =
0.001
timediff =
0.009
R2 =
0.190782119898936
R2_p =
0.001
---
0.0 3.0
coefficients :
Intercept =
-3.0200148063588657
cyc =
0.8724460094246678
timediff =
-0.0029619890630369398
coefficient_pvals :
Intercept =
0.988
cyc =
0.001
timediff =
0.022
R2 =
0.15683570137446723
R2_p =
0.001
---
0.0 3.5
coefficients :
Intercept =
-3.2295762867290967
cyc =
0.9149179573694931
timediff =
-0.0026384195441538417
coefficient_pvals :
Intercept =
0.986
cyc =
0.001
timediff =
0.022
R2 =
0.15072357787385882
R2_p =
0.001
---
```

```
0.5 0.0
coefficients :
Intercept =
-0.8491182889162299
cyc =
0.08838652915315819
timediff =
-0.00043990170681410077
coefficient_pvals :
Intercept =
0.718
cyc =
0.09
timediff =
0.581
R2 =
0.009239244556658144
R2_p =
0.487
---
0.5 0.5
coefficients :
Intercept =
-0.5218286238299308
cyc =
0.13416461139906222
timediff =
-0.0006463105262043438
coefficient_pvals :
Intercept =
0.785
cyc =
0.012
timediff =
0.434
R2 =
0.017147628673866544
R2_p =
0.303
---
0.5 1.0
coefficients :
Intercept =
1.2324676591848363
cyc =
0.25141066227936626
timediff =
-0.0013030509016013307
coefficient_pvals :
Intercept =
0.076
cyc =
0.002
timediff =
0.197
R2 =
0.04553576382920732
R2_p =
0.085
---
0.5 1.5
coefficients :
Intercept =
6.5265894635452755
cyc =
0.5735312928109829
timediff =
-0.0028002399876733298
coefficient_pvals :
Intercept =
0.039
cyc =
0.001
timediff =
0.087
R2 =
0.07186934066793693
R2_p =
0.014
---
```

```
0.5 2.0
coefficients :
Intercept =
5.735245593506558
cyc =
0.7755169628258052
timediff =
-0.0025929447483702206
coefficient_pvals :
Intercept =
0.067
cyc =
0.001
timediff =
0.16
R2 =
0.07987737922470672
R2_p =
0.015
---
0.5 2.5
coefficients :
Intercept =
-2.567297402467069
cyc =
0.7252972132561898
timediff =
-0.0025145208698113273
coefficient_pvals :
Intercept =
0.993
cyc =
0.001
timediff =
0.015
R2 =
0.1755481943178887
R2_p =
0.001
---
0.5 3.0
coefficients :
Intercept =
-4.379004766873002
cyc =
0.5138637162318845
timediff =
-0.001987416940390203
coefficient_pvals :
Intercept =
0.98
cyc =
0.005
timediff =
0.047
R2 =
0.09387224679016559
R2_p =
0.004
---
0.5 3.5
coefficients :
Intercept =
-4.5795867684094835
cyc =
0.4922739989294412
timediff =
-0.0016549883402800327
coefficient_pvals :
Intercept =
0.917
cyc =
0.008
timediff =
0.149
R2 =
0.06151091890663285
R2_p =
0.019
---
```

```
1.0 0.0
coefficients :
Intercept =
-0.6656069451529981
cyc =
0.04830405484756581
timediff =
-0.00013850519931090649
coefficient_pvals :
Intercept =
0.697
cyc =
0.026
timediff =
0.599
R2 =
0.012558526261000957
R2_p =
0.381
---
1.0 0.5
coefficients :
Intercept =
-0.3185223858210026
cyc =
0.05812531508652023
timediff =
-0.00022776158408926276
coefficient_pvals :
Intercept =
0.789
cyc =
0.025
timediff =
0.439
R2 =
0.020888405245899966
R2_p =
0.277
---
1.0 1.0
coefficients :
Intercept =
1.1610818157113516
cyc =
0.11146115307238548
timediff =
-0.0005315441383797039
coefficient_pvals :
Intercept =
0.069
cyc =
0.004
timediff =
0.164
R2 =
0.0523818394308585
R2_p =
0.053
---
1.0 1.5
coefficients :
Intercept =
5.680956578114451
cyc =
0.41760598652730435
timediff =
-0.0012318244806533804
coefficient_pvals :
Intercept =
0.083
cyc =
0.001
timediff =
0.184
R2 =
0.08413553637140803
R2_p =
0.009
---
```

```
1.0 2.0
coefficients :
Intercept =
3.4948743092481585
cyc =
0.17885948477205835
timediff =
-0.00022842939818943693
coefficient_pvals :
Intercept =
0.339
cyc =
0.003
timediff =
0.757
R2 =
0.016039380397784186
R2_p =
0.307
---
1.0 2.5
coefficients :
Intercept =
-1.9351892100210852
cyc =
-0.08813814498481463
timediff =
-0.0002731641628889695
coefficient_pvals :
Intercept =
0.727
cyc =
0.093
timediff =
0.522
R2 =
0.011912002746543693
R2_p =
0.373
---
1.0 3.0
coefficients :
Intercept =
-3.8686697797830316
cyc =
-0.3058510153161406
timediff =
-0.00033371343509584436
coefficient_pvals :
Intercept =
0.655
cyc =
0.002
timediff =
0.61
R2 =
0.045554493460762124
R2_p =
0.043
---
1.0 3.5
coefficients :
Intercept =
-4.868830678054428
cyc =
-0.4851549267270315
timediff =
0.0005812773212846924
coefficient_pvals :
Intercept =
0.183
cyc =
0.001
timediff =
0.442
R2 =
0.07306014425279983
R2_p =
0.008
---
```

```
1.5 0.0
coefficients :
Intercept =
-0.5011485782662916
cyc =
-0.00633730844116448
timediff =
-0.00024076056214923628
coefficient_pvals :
Intercept =
0.906
cyc =
0.69
timediff =
0.202
R2 =
0.02272201968042009
R2_p =
0.221
---
1.5 0.5
coefficients :
Intercept =
-0.2542492189015602
cyc =
-0.00873810772109919
timediff =
-0.00030931084478127936
coefficient_pvals :
Intercept =
0.939
cyc =
0.609
timediff =
0.105
R2 =
0.03541933655838514
R2_p =
0.113
---
1.5 1.0
coefficients :
Intercept =
0.8081220098054908
cyc =
0.039050717366603524
timediff =
-0.0004419222739826697
coefficient_pvals :
Intercept =
0.026
cyc =
0.161
timediff =
0.046
R2 =
0.04904582280716119
R2_p =
0.032
---
1.5 1.5
coefficients :
Intercept =
3.5716205908405136
cyc =
0.30066690080115477
timediff =
-0.00029145143862434054
coefficient_pvals :
Intercept =
0.34
cyc =
0.001
timediff =
0.717
R2 =
0.04081697624634473
R2_p =
0.086
---
```

```
1.5 2.0
coefficients :
Intercept =
1.8623819694354966
cyc =
0.12062036276062588
timediff =
0.0009228463183984592
coefficient_pvals :
Intercept =
0.953
cyc =
0.032
timediff =
0.1
R2 =
0.03997516451050309
R2_p =
0.078
---
1.5 2.5
coefficients :
Intercept =
-0.8010410812692712
cyc =
-0.12682374800142834
timediff =
0.0002872091580009348
coefficient_pvals :
Intercept =
0.162
cyc =
0.015
timediff =
0.366
R2 =
0.03165787397468611
R2_p =
0.081
---
1.5 3.0
coefficients :
Intercept =
-2.0283888972193664
cyc =
-0.5352989730134395
timediff =
-0.0004352664075232857
coefficient_pvals :
Intercept =
0.7
cyc =
0.001
timediff =
0.487
R2 =
0.12682988730915223
R2_p =
0.001
---
1.5 3.5
coefficients :
Intercept =
-3.3410915947549777
cyc =
-0.7634198957192057
timediff =
0.0005567074842592311
coefficient_pvals :
Intercept =
0.189
cyc =
0.001
timediff =
0.453
R2 =
0.19069093926484215
R2_p =
0.001
---
```

```
2.0 0.0
coefficients :
Intercept =
-0.4217182582864356
cyc =
-0.0323147834645035
timediff =
-0.00042188425654563096
coefficient_pvals :
Intercept =
0.874
cyc =
0.253
timediff =
0.27
R2 =
0.02077843069838381
R2_p =
0.273
---
2.0 0.5
coefficients :
Intercept =
-0.26119737753838074
cyc =
-0.04109341166572737
timediff =
-0.000501258680453274
coefficient_pvals :
Intercept =
0.905
cyc =
0.209
timediff =
0.166
R2 =
0.027007814950561038
R2_p =
0.171
---
2.0 1.0
coefficients :
Intercept =
0.42803873259300007
cyc =
-0.050601377267051065
timediff =
-0.0005307696560791606
coefficient_pvals :
Intercept =
0.018
cyc =
0.117
timediff =
0.032
R2 =
0.04822528172241258
R2_p =
0.03
---
2.0 1.5
coefficients :
Intercept =
2.1292322827050194
cyc =
0.0038036941242172227
timediff =
-6.380406970809368e-05
coefficient_pvals :
Intercept =
0.443
cyc =
0.949
timediff =
0.922
R2 =
0.00012227822439359048
R2_p =
0.984
---
```

```
2.0 2.0
coefficients :
Intercept =
1.1245191387015383
cyc =
0.08444619974278715
timediff =
0.0012653401458255853
coefficient_pvals :
Intercept =
0.991
cyc =
0.183
timediff =
0.026
R2 =
0.055466821920785
R2_p =
0.024
---
2.0 2.5
coefficients :
Intercept =
-0.46807479857920253
cyc =
0.15994810876384657
timediff =
0.0007264792446811663
coefficient_pvals :
Intercept =
0.019
cyc =
0.008
timediff =
0.041
R2 =
0.056964638169029524
R2_p =
0.016
---
2.0 3.0
coefficients :
Intercept =
-0.7949340233973721
cyc =
-0.3140551453889507
timediff =
-0.0006544727215219011
coefficient_pvals :
Intercept =
0.872
cyc =
0.001
timediff =
0.227
R2 =
0.06856246373260866
R2_p =
0.01
---
2.0 3.5
coefficients :
Intercept =
-1.6817608523347396
cyc =
-0.552696722970635
timediff =
-5.8531652433809905e-05
coefficient_pvals :
Intercept =
0.467
cyc =
0.001
timediff =
0.925
R2 =
0.11883864226937901
R2_p =
0.002
---
```

```
2.5 0.0
coefficients :
Intercept =
-0.3362677443307011
cyc =
-0.046500459327780214
timediff =
-0.0005823489292851276
coefficient_pvals :
Intercept =
0.853
cyc =
0.255
timediff =
0.301
R2 =
0.01821246383703734
R2_p =
0.313
---
2.5 0.5
coefficients :
Intercept =
-0.23138415386916159
cyc =
-0.0564158745451604
timediff =
-0.000659612106709769
coefficient_pvals :
Intercept =
0.898
cyc =
0.205
timediff =
0.214
R2 =
0.021153616279858678
R2_p =
0.218
---
2.5 1.0
coefficients :
Intercept =
0.19159053384358374
cyc =
-0.10111499140536527
timediff =
-0.0006297667700496915
coefficient_pvals :
Intercept =
0.062
cyc =
0.04
timediff =
0.109
R2 =
0.0382610201084137
R2_p =
0.071
---
2.5 1.5
coefficients :
Intercept =
1.357692336802497
cyc =
-0.19728400754750963
timediff =
-0.00023640508117473214
coefficient_pvals :
Intercept =
0.37
cyc =
0.018
timediff =
0.71
R2 =
0.021055272569028838
R2_p =
0.186
---
```

```
2.5 2.0
coefficients :
Intercept =
0.9260130887649884
cyc =
-0.043850594896681826
timediff =
0.001082552316264891
coefficient_pvals :
Intercept =
0.962
cyc =
0.467
timediff =
0.075
R2 =
0.03883469091903313
R2_p =
0.071
---
2.5 2.5
coefficients :
Intercept =
-0.49298662762254736
cyc =
0.3063885589517678
timediff =
0.0011562461940411192
coefficient_pvals :
Intercept =
0.006
cyc =
0.002
timediff =
0.04
R2 =
0.08615724015510295
R2_p =
0.009
---
2.5 3.0
coefficients :
Intercept =
-0.32789624678340323
cyc =
-4.510571545370318e-05
timediff =
-0.0004538599286528144
coefficient_pvals :
Intercept =
0.888
cyc =
0.999
timediff =
0.223
R2 =
0.012270834444229606
R2_p =
0.383
---
2.5 3.5
coefficients :
Intercept =
-0.6892975772766514
cyc =
-0.26772477019492
timediff =
-0.00043618469790457565
coefficient_pvals :
Intercept =
0.754
cyc =
0.002
timediff =
0.421
R2 =
0.04603539309161564
R2_p =
0.033
---
```

```
3.0 0.0
coefficients :
Intercept =
-0.2557868519656272
cyc =
-0.06139484132764147
timediff =
-0.0006914142271712679
coefficient_pvals :
Intercept =
0.842
cyc =
0.237
timediff =
0.29
R2 =
0.01732745868754959
R2_p =
0.288
---
3.0 0.5
coefficients :
Intercept =
-0.18025631628306582
cyc =
-0.06910384717350801
timediff =
-0.0007542705469859821
coefficient_pvals :
Intercept =
0.862
cyc =
0.192
timediff =
0.266
R2 =
0.018590825520710563
R2_p =
0.263
---
3.0 1.0
coefficients :
Intercept =
0.09168324197638962
cyc =
-0.09993829305310961
timediff =
-0.0006906401424738534
coefficient_pvals :
Intercept =
0.527
cyc =
0.104
timediff =
0.204
R2 =
0.025362090822878103
R2_p =
0.178
---
3.0 1.5
coefficients :
Intercept =
0.9376116583625811
cyc =
-0.2684434287020205
timediff =
-0.0004375602361984047
coefficient_pvals :
Intercept =
0.208
cyc =
0.008
timediff =
0.373
R2 =
0.04300489567502597
R2_p =
0.029
---
```

```
3.0 2.0
coefficients :
Intercept =
0.9103659653528108
cyc =
-0.19257307967061876
timediff =
0.0006027663155113364
coefficient_pvals :
Intercept =
0.836
cyc =
0.01
timediff =
0.354
R2 =
0.0327619948233534
R2_p =
0.105
---
3.0 2.5
coefficients :
Intercept =
-0.4449087268556247
cyc =
0.26417310272561056
timediff =
0.0014204332774473193
coefficient_pvals :
Intercept =
0.006
cyc =
0.012
timediff =
0.025
R2 =
0.07185138844636973
R2_p =
0.014
---
3.0 3.0
coefficients :
Intercept =
-0.2917801899380713
cyc =
0.3150475184579608
timediff =
-0.00019408401427533912
coefficient_pvals :
Intercept =
0.769
cyc =
0.001
timediff =
0.554
R2 =
0.09849941701340803
R2_p =
0.003
---
3.0 3.5
coefficients :
Intercept =
-0.20636932318703635
cyc =
-0.09156639908883314
timediff =
-0.0005649954991944523
coefficient_pvals :
Intercept =
0.872
cyc =
0.161
timediff =
0.217
R2 =
0.020407347263527886
R2_p =
0.198
---
```

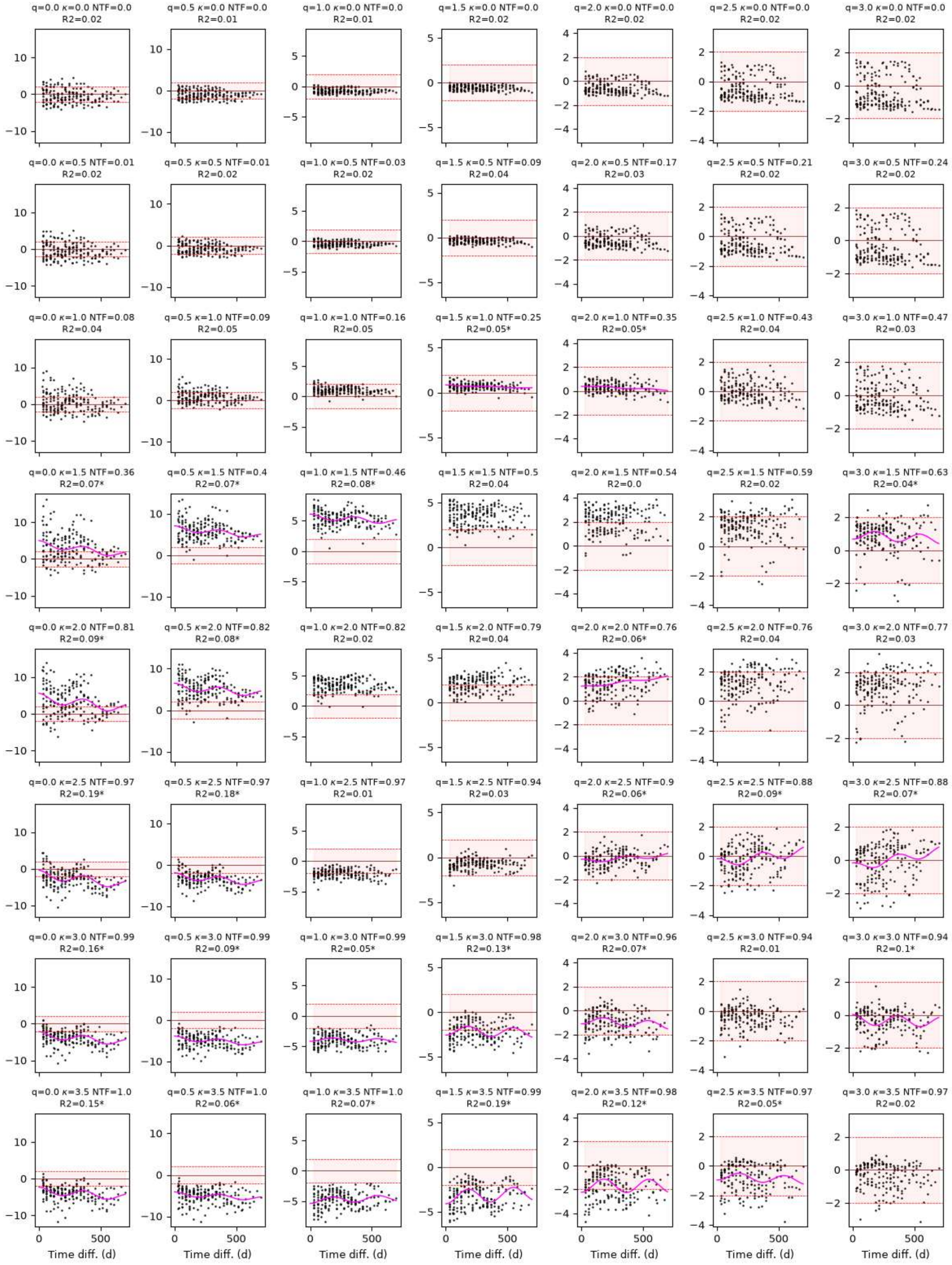


### Plot figure 6

In [14]:
```
import qdiv
import pandas as pd
import pickle
```

```python
import numpy as np
import math
import matplotlib.pyplot as plt
import matplotlib.dates as mdates
import matplotlib.gridspec as gridspec
from matplotlib.colors import LinearSegmentedColormap
from matplotlib.colors import BoundaryNorm
path = 'case_MBR_folder/'

with open(path + 'Stats_beta.pickle', 'rb') as f:
    data = pickle.load(f)

colors = ['#bdbdbd', '#969696', '#636363', '#252525']
qlist = [0.0, 0.5, 1.0, 1.5, 2.0, 2.5, 3.0]
loclist = [0.0, 0.5, 1.0, 1.5, 2.0, 2.5, 3.0, 3.5]

ses_mean = pd.DataFrame(pd.NA, index=loclist, columns=qlist)
ses_std = pd.DataFrame(pd.NA, index=loclist, columns=qlist)
ntf_mean = pd.DataFrame(pd.NA, index=loclist, columns=qlist)
ntf_std = pd.DataFrame(pd.NA, index=loclist, columns=qlist)
R2 = pd.DataFrame(pd.NA, index=loclist, columns=qlist)
R2p = pd.DataFrame(pd.NA, index=loclist, columns=qlist)
cycp = pd.DataFrame(pd.NA, index=loclist, columns=qlist)

for k in data.keys():
    loc = data[k]['loc']
    q = data[k]['q']
    ses_mean.loc[loc, q] = data[k]['ses'].mean()
    ses_std.loc[loc, q] = data[k]['ses'].std()
    ntf_mean.loc[loc, q] = data[k]['NTF'].mean()
    ntf_std.loc[loc, q] = data[k]['NTF'].std()
    R2.loc[loc, q] = data[k]['R2']
    R2p.loc[loc, q] = data[k]['R2_p']
    cycp.loc[loc, q] = data[k]['cyc_p']

plt.rcParams.update({'font.size':10})
fig = plt.figure(figsize=(18/2.54, 13/2.54))
fig.subplots_adjust(wspace=0.5, hspace=0.7)
gs = gridspec.GridSpec(2, 6, height_ratios=[2, 1])
ax1 = fig.add_subplot(gs[0,:3])
ax2 = fig.add_subplot(gs[0,3:])
ax3 = fig.add_subplot(gs[1,:2])
ax4 = fig.add_subplot(gs[1,2:4])
ax5 = fig.add_subplot(gs[1,4:])

ntf = np.asarray(ntf_mean.T.to_numpy(), dtype=float)

# Heatmap qiNRI mean values
bounds = [-6, -2, 0, 2, 6]
cmap = LinearSegmentedColormap.from_list(
    'ses',
    [
        '#08306B',  # extreme negative
        '#2166AC',  # -2
        '#FFFFFF',  # -0.5
        '#FFFFFF',  # +0.5
        '#B2182B',  # +2
        '#67000D'   # extreme positive
    ],
    N=256
)
bounds = [-6, -2, -0.2, 0.2, 2, 6]
norm = BoundaryNorm(bounds, cmap.N)

df = ses_mean
X, Y = np.meshgrid(df.index, df.columns)
Zvals = np.asarray(df.T.to_numpy(), dtype=float)
zmin = Zvals.min(); zmax = Zvals.max()

cf = ax1.contourf(
    X, Y, Zvals,
    levels=np.linspace(-6, 6, 101),
    cmap=cmap,
    norm=norm
)
cs = ax1.contour(X, Y, ntf, levels=[0.2, 0.5, 0.7, 0.9, 0.97], colors='lightgrey', linewidths=0.5)
ax1.clabel(cs, fontsize=7) #Contour labels
cbar = fig.colorbar(cf, ax=ax1, location='top')
cbar.set_ticks([-4, -2, 0, 2, 4])
cbar.set_label(r'(a) Mean $\beta$-$^q$iNRI', fontsize=9)
```

```
ax1.set_yticks(range(4))
ax1.set_ylabel('q')
ax1.set_xticks(range(4))
ax1.set_xlabel(r'$\kappa$')

# Heatmap R2 values for significant fits
df = R2.copy()
#df[cycp>0.05] = np.nan
cmap = plt.cm.viridis.copy()
cmap.set_bad('white')
X, Y = np.meshgrid(df.index, df.columns)
Zvals = np.asarray(df.T.to_numpy(), dtype=float)
zmin = Zvals[np.isfinite(Zvals)].min(); zmax = Zvals[np.isfinite(Zvals)].max()

cf = ax2.contourf(
    X, Y, Zvals,
    levels=np.linspace(0, 0.2, 101),
    cmap=cmap,
    vmin=0, vmax=0.2,
)

# grey overlay for non-significant regions
mask = (R2p.T.to_numpy() > 0.05)
ax2.contourf(
    X, Y, mask.astype(float),
    levels=[0.5, 1.5],
    colors=['lightgrey'],
    alpha=0.4
)

cs = ax2.contour(X, Y, ntf, levels=[0.2, 0.5, 0.7, 0.9, 0.97],
                 colors='white', linewidths=0.5
                )
ax2.clabel(cs, fontsize=7) #Contour labels

cbar = fig.colorbar(cf, ax=ax2, location='top')
cbar.set_ticks([0, 0.1, 0.2])
cbar.set_label('(b) R$^2$ (p<0.05)', fontsize=9)
ax2.set_yticks(range(4))
ax2.set_yticklabels([])
ax2.set_xticks(range(4))
ax2.set_xlabel(r'$\kappa$')

for i, (ax, key) in enumerate(zip([ax3, ax4, ax5],
                                  ['q=1.0_loc=0.5', 'q=1.0_loc=1.5', 'q=1.0_loc=3.0'])
                             ):
    x = data[key]['x']
    ses = data[key]['ses']
    ax.scatter(x, ses, marker='.', color='grey', s=2, alpha=0.5)
    ax.axhline(-2, 0, 1, color='red', lw=0.5, ls='--')
    ax.axhline(0, 0, 1, color='darkred', lw=0.5, ls='-')
    ax.axhline(2, 0, 1, color='red', lw=0.5, ls='--')
    ax.fill_between([0, 700], -2, 2, color='red', alpha=0.05)
    if 'xrange' in data[key]:
        ax.plot(data[key]['xrange'], data[key]['yrange'], color='black', lw=1)
    ax.set_ylim(-6, 8)
    ax.set_yticks(range(-6, 8, 3))
    if i != 0:
        ax.set_yticklabels([])
    else:
        ax.set_ylabel(r'$\beta$-$^q$iNRI')
    ax.set_xlim(0, 700)
    ax.set_xticks(range(0, 700, 200))
    ax.set_xlabel('Time diff. (d)')

    q = str(data[key]['q'])
    loc = str(data[key]['loc'])
    ntf = str(round(data[key]['NTF'].mean(), 2))
    title = r'('+list('cde')[i]+')'+' q=' + q + ' $\kappa$=' + loc + '\nNTF=' + ntf
    ax.set_title(title, fontsize=9)

    for spine in ['top', 'right']:
        ax.spines[spine].set_visible(False)
    for spine in ['left', 'bottom']:
        ax.spines[spine].set_linewidth(0.8)

    ax1.text(float(loc), float(q), list('cde')[i], va='center', ha='center', color='lightgrey')
    ax2.text(float(loc), float(q), list('cde')[i], va='center', ha='center', color='white')
```

```
plt.savefig(path + 'Fig_beta_iNRIq_heatmap_and_vs_sampling_time', dpi=600)
plt.savefig(path + 'Fig_beta_iNRIq_heatmap_and_vs_sampling_time.pdf', format='pdf')
```

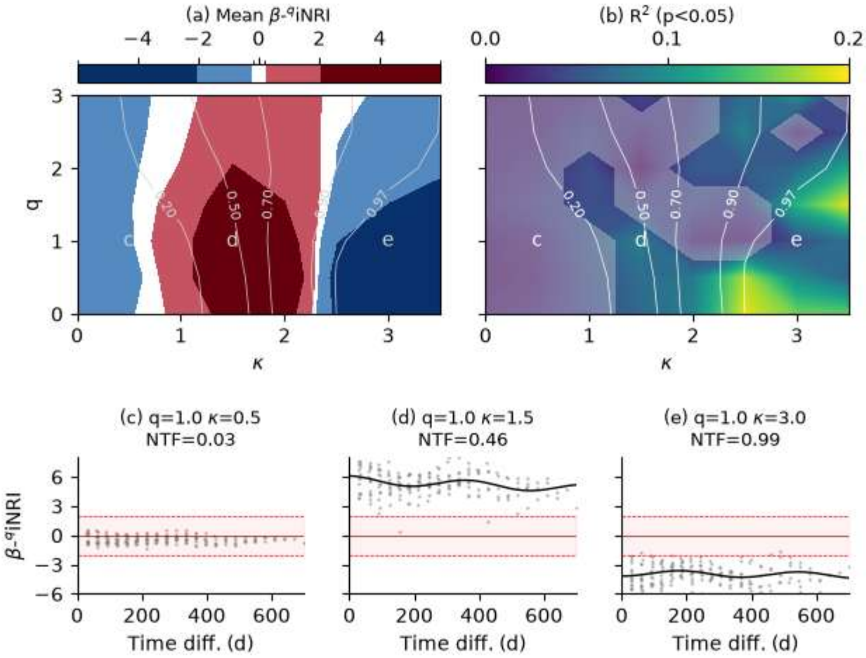
